# Local albumin excess exacerbates Candida albicans-induced inflammasome activation linked with hyperinflammation during vulvovaginal candidiasis

**DOI:** 10.64898/2026.01.22.700771

**Authors:** Sophie Austermeier, Axel Dietschmann, Gianluca Vascelli, Kar On Cheng, Sophia U. J. Hitzler, Kira S. Skurk, Stefanie Westermann, Dolly E. Montaño, Martin Jaeger, Nadja Jablonowski, Stephanie Wisgott, Osama Elshafee, Chiara Brachelente, Dennis M. de Graaf, Tomasz Próchnicki, Eicke Latz, Stefan Wirtz, Christoph Becker, Giorgia Renga, Vasileios Oikonomou, Bernhard Hube, Gilbert G.G. Donders, Marina Pekmezovic, Teresa Zelante, Mark S. Gresnigt

## Abstract

Vulvovaginal candidiasis (VVC) is a mucosal yeast infection where symptoms are driven by inflammatory responses. The onset of VVC is instigated by the yeast *Candida albicans*, but the underlying causes of disease-driving hyperinflammation remains incompletely resolved. We found that vaginal albumin concentrations are higher in women with recurrent VVC and during VVC in mice. These albumin levels correlated with inflammatory cytokines in vaginal lavages. While the abundance of this serum protein in the vagina can represent a hallmark of inflammation-induced vascular permeability, we reveal molecular mechanisms by which albumin also drives inflammation. It increased NLRP3 inflammasome activation in human macrophages elicited by *C. albicans*. Albumin induced fungal adaptations that increased resistance to macrophage-mediated clearance. Further it increased the expression of fungal secreted aspartic protease 1, which we identified as the effector driving inflammasome activation. Moreover, neutrophils show impaired effectivity in clearing *C. albicans* in presence of albumin. Collectively, vaginal albumin may not only be a consequence of inflammation of the vaginal mucosa, but also can drive hyperinflammatory responses that underlie immunopathology in VVC.

**Graphical abstract:** 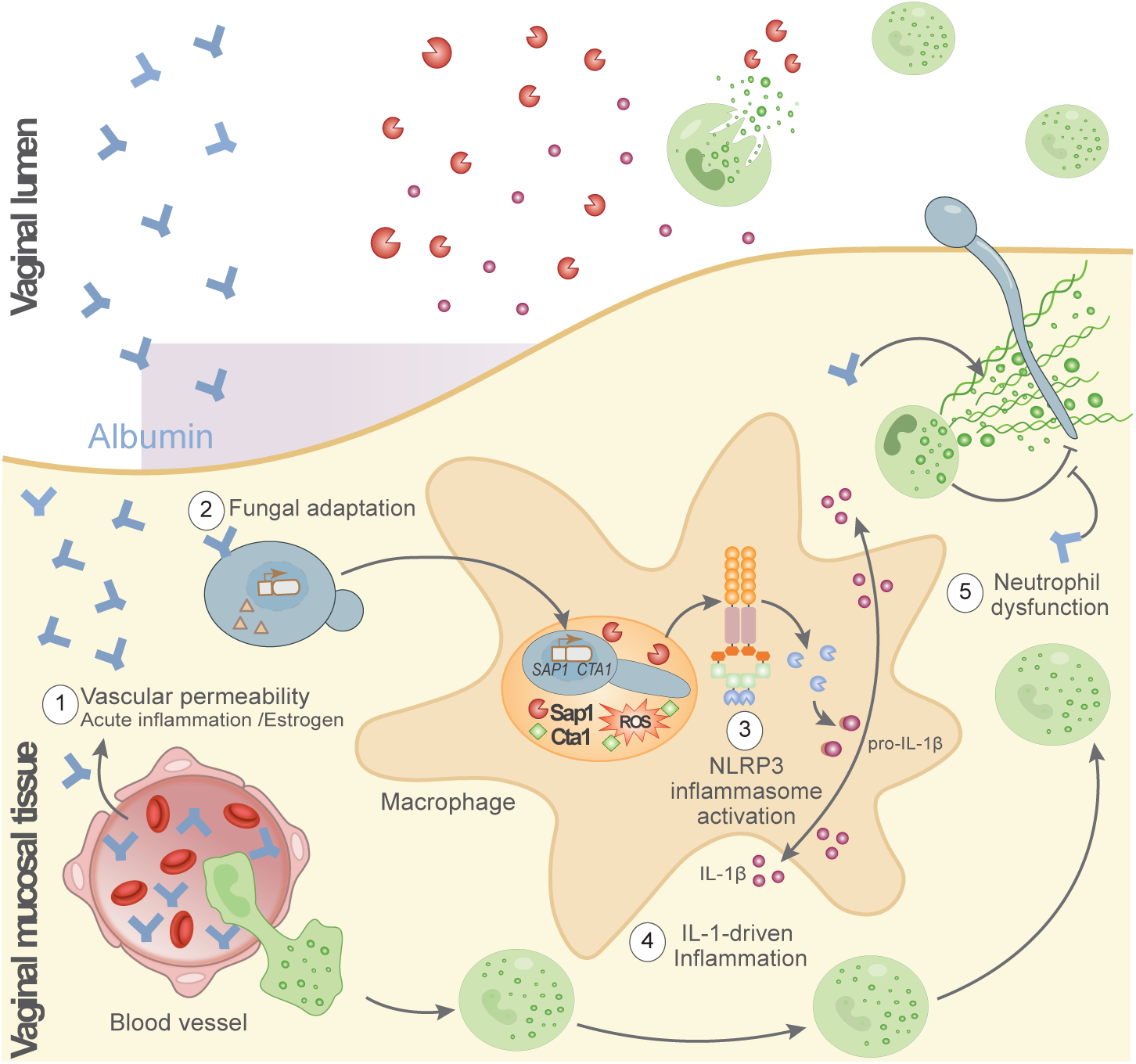

## INTRODUCTION

Vulvovaginal candidiasis (VVC) is one of the most common infections of the female reproductive tract, occurring in 70-75% of otherwise healthy women at least once during their life (1). Particularly for women experiencing several episodes per year, referred to as recurrent VVC (RVVC), quality of life is severely affected (2, 3). VVC symptoms are driven by an inflammatory response against the mucosal infection with *Candida* species (4), most commonly *C. albicans* (5). This inflammatory response is instigated by fungal pathogenicity, though its persistence and development of symptoms result from persistent neutrophil responses, in part driven by activation of the NLRP3 inflammasome (4, 6–8). The recruited neutrophils during VVC are dysfunctional in fungal clearance and cause immunopathology to the vulvovaginal mucosa (6, 8).

The pathogenicity of *C. albicans* in the context of VVC involves several processes. The toxin candidalysin causes epithelial damage and triggers immune responses in the vaginal mucosa that drive downstream inflammation (9–11). The trace mineral binding protein Pra1 helps *C. albicans* to overcome zinc starvation, yet can be recognized by the immune system to elicit neutrophil recruitment (12). Other infection-specific proteins known to be involved during VVC are *C. albicans* secreted aspartic proteinases (Saps) (13). Some of these virulence factors trigger the activation of the NLRP3 inflammasome (14, 15), a multiprotein complex that is highly expressed in macrophages (16), which are abundant immune cells in both the human and murine vaginal mucosa (17–20). This inflammasome processes pro-interleukin (IL)-1β into its bioactive proinflammatory form (21), which subsequently is a strong driver of neutrophil-mediated inflammation during VVC (7, 22, 23). While the association between NLRP3 inflammasome activation and VVC is clear, it is not completely understood what drives its characteristic hyperactivation in this disease.

VVC mainly affects women in their fertile years (24). Among other predisposing factors, the use of hormonal replacement therapy, pregnancy, and the use of oral contraceptives are associated with VVC (1, 24, 25). During the fertile years, estradiol is the predominant form of estrogen in the female body and its production depends on the menstrual cycle (26). To mimic and study VVC using mouse models, estradiol administration is required to successfully induce *C. albicans* infections in the vaginal tract. Collectively, this highlights the central role of estradiol in the susceptibility to VVC. Besides its major role in reproductive health, estradiol increases the vascular permeability of the female reproductive organs (27). Acute local inflammatory responses can further increase the vascular permeability. Consequently, albumin, as the most abundant serum protein, is prevalently found in the vaginal fluid, particularly in the context of inflammation markers (28). We previously identified that adaptations of *C. albicans* and *C. glabrata* in response to albumin can enhance their pathogenicity towards host epithelial cells (29, 30), yet albumin can also neutralize candidalysin (31), a key driver of VVC-associated inflammation and inflammasome activation(7, 15, 22, 23). Thus, while is present in the vagina during inflammation, its role in the inflammatory pathogenesis of VVC remains unclear.

Here, we demonstrate that vaginal albumin levels not only significantly correlate with inflammation during experimental VVC in mice and in patients with recurrent VVC, but also that albumin is a mechanistic driver of *C. albicans*-induced hyperinflammatory responses. Albumin was observed to induce *C. albicans* pathogenicity mechanisms to subsequently elicit stronger NLRP3 inflammasome activation.

## RESULTS

### Vaginal albumin levels are linked to inflammation during VVC

To define the association of vaginal albumin with VVC, albumin was quantified in vaginal lavages from a previously collected cohort of healthy women and RVVC patients (32). In line with previous observations (28), vaginal albumin concentrations were elevated in RVVC patients compared to healthy women (**Figure 1A**). To understand how albumin is linked with the inflammatory response during RVVC, a Pearson correlation analysis was performed. Significant correlations between albumin concentrations and the proinflammatory cytokines IL-1β, IL-6, and IL-17, and the neutrophil chemoattractant IL-8, were found in vaginal lavages from RVVC patients (RVVC_1_), but not in healthy women (**Figure 1 B+C**, **Supplementary Figure S1**). To validate this link in RVVC patients, samples from an independent multi-center patient cohort (RVVC_2_) were analyzed (33, 34). We similarly observed significant correlations of IL-1β, IL-6 and IL-8 with the abundance of albumin. Additionally, IL-1 receptor antagonist (Ra) and TNF positively correlated to albumin concentrations in the vaginal lavages collected within the RVVC_2_ cohort (**Figure 1B, Supplementary Figure S1**). To validate the link between albumin and parameters of VVC immunopathology, we performed an estradiol-based mouse VVC model (**Figure 1D**). We observed establishment of fungal invasion and a strong neutrophilic cell infiltration into vaginal tissue and lumen (**Figure 1E**). Further linking albumin and inflammation during VVC, we found albumin concentrations significantly increased in vaginal lavages and tissue of *C. albicans*-infected β-estradiol-primed mice (**Figure 1F**). One-week post infection, vaginal albumin correlated with features of the inflammatory response of VVC such as vaginal IL-1β concentrations and myeloperoxidase (MPO) levels, together hinting at neutrophilic inflammation. Further, a trend (*P* = 0.09) towards increased neutrophil numbers in week 1. A significant correlation with neutrophil numbers in week 2 with albumin levels suggest that stronger inflammatory responses early during infection drive persistent neutrophilia (**Figure 1G**). Two weeks post infection, a positive correlation between albumin and MPO release suggested a persistent link albumin and neutrophil activation later during infection (**Supplementary Figure S2**).

**Figure 1:**
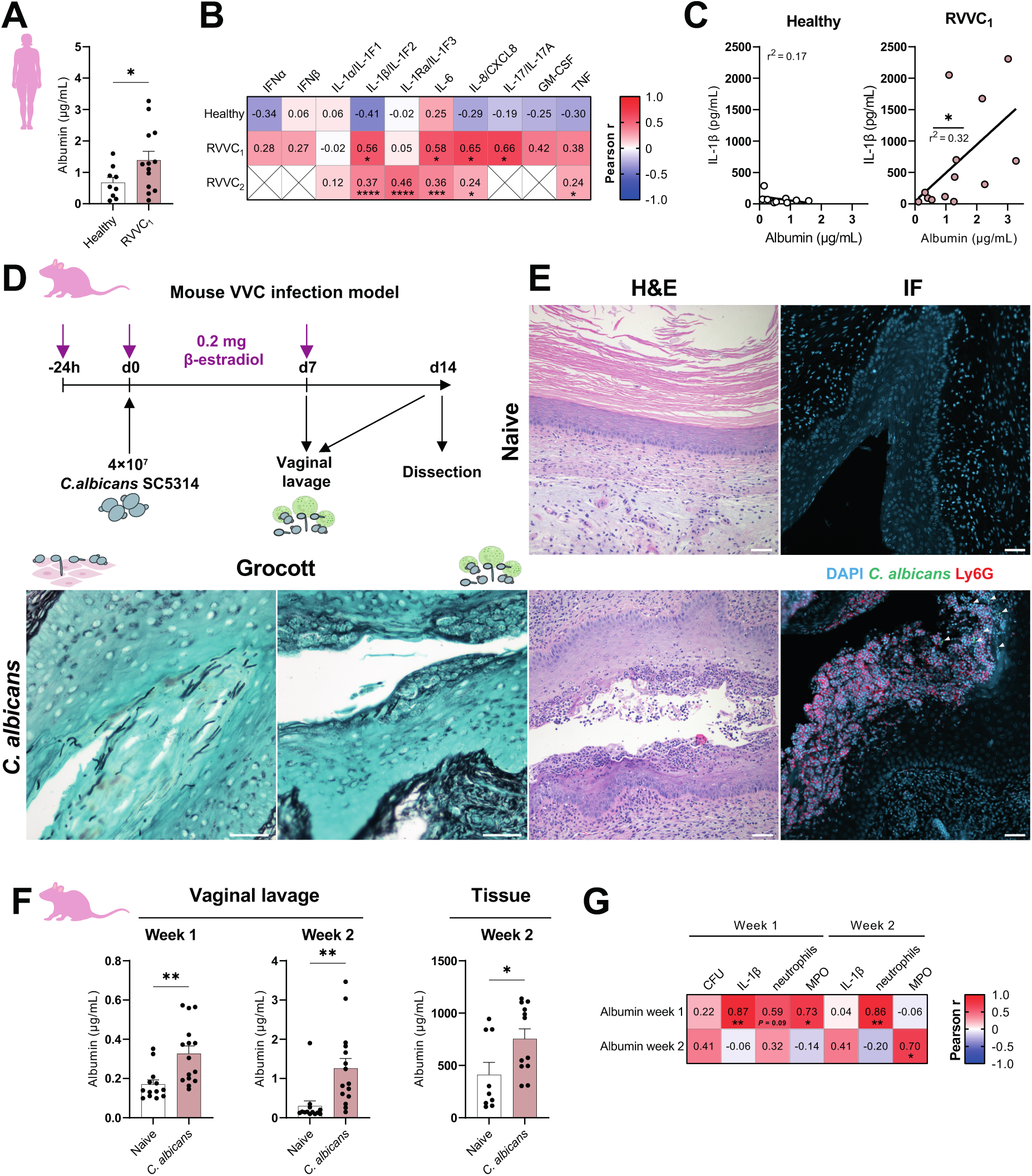
Albumin concentrations correlate with VVC-associated cytokines *in vivo*. **(A)** Albumin concentrations in vaginal lavages of healthy women (*n* = 9) and RVVC_1_ patients (*n* = 13). Significance was tested using a Welch’s t-test. **(B)** Heatmap showing Pearson correlation coefficient (r^2^) of cytokine and albumin concentrations in vaginal lavages from healthy women (*n* = 9) and two independent cohorts of RVVC patients (*n* = 13 and *n* = 129 respectively). **(C)** Example of correlations shown in (B) for IL-1β and albumin in the RVVC_1_ cohort and healthy women. **(D)** Mouse VVC infection model scheme. **(E)** Representative images of vaginal histology sections of naïve or *C. albicans* treated mice, 2 weeks after infection. Sections were stained with hematoxylin/eosin to visualize tissue pathology (H&E, scale = 100 µm), Grocott’s (Scale = 50 µm) to visualize fungal presence and immunofluorescence (scale = 50 µm) showing nuclei (DAPI, blue), *C. albicans* (anti-*Candida* antibody, green, indicated by white arrowheads) and neutrophils (anti-Ly6G, red). Images are representative of 4 mice per group. **(F)** Albumin levels in vaginal lavages (*n* = 13 uninfected and *n =* 15 infected) or organ homogenate (*n* = 9 uninfected and *n* = 12 infected) from β-estradiol-treated, and β-estradiol-treated *C. albicans*-infected mice. Significances were calculated using Welch’s t-test. **(G)** Pearson correlations between albumin concentrations and colony forming units (CFUs, week 1), IL-1β concentrations, MPO concentrations (*n* = 9) or neutrophil count (*n* = 9) in vaginal lavages one week after infection of β-estradiol-treated mice with *C. albicans.* Data in F and G are pooled data from three and two independently conducted experiments respectively. Bars represent the mean + SEM with dots as individual biological replicates. * = *P ≤* 0.05, ** = *P ≤* 0.01, *** = *P ≤* 0.001, **** = *P ≤* 0.0001

### Albumin potentiates innate inflammatory responses

Epithelial cells mount initial inflammatory responses to vaginal *C. albicans* infection and recruit and activate neutrophils (7, 10). To dissect whether albumin may not only be a consequence but also a driver of inflammation, we introduced albumin to an *in vitro* vulvovaginal epithelial infection model (A-431 cells). The presence of albumin did not affect *C. albicans*-driven IL-1α and IL-1Ra responses and even decreased IL-8 release by A-431 cells (**Figure 2A**). Moreover, the cytokines IL-1β and IL-6 were not released by these cells (**Figure 2A**). Even following prolonged incubation, a setting where albumin increases *C. albicans-*induced epithelial damage (30), no substantial increase in the release of proinflammatory cytokines was observed (**Supplementary Figure S3**).

**Figure 2:**
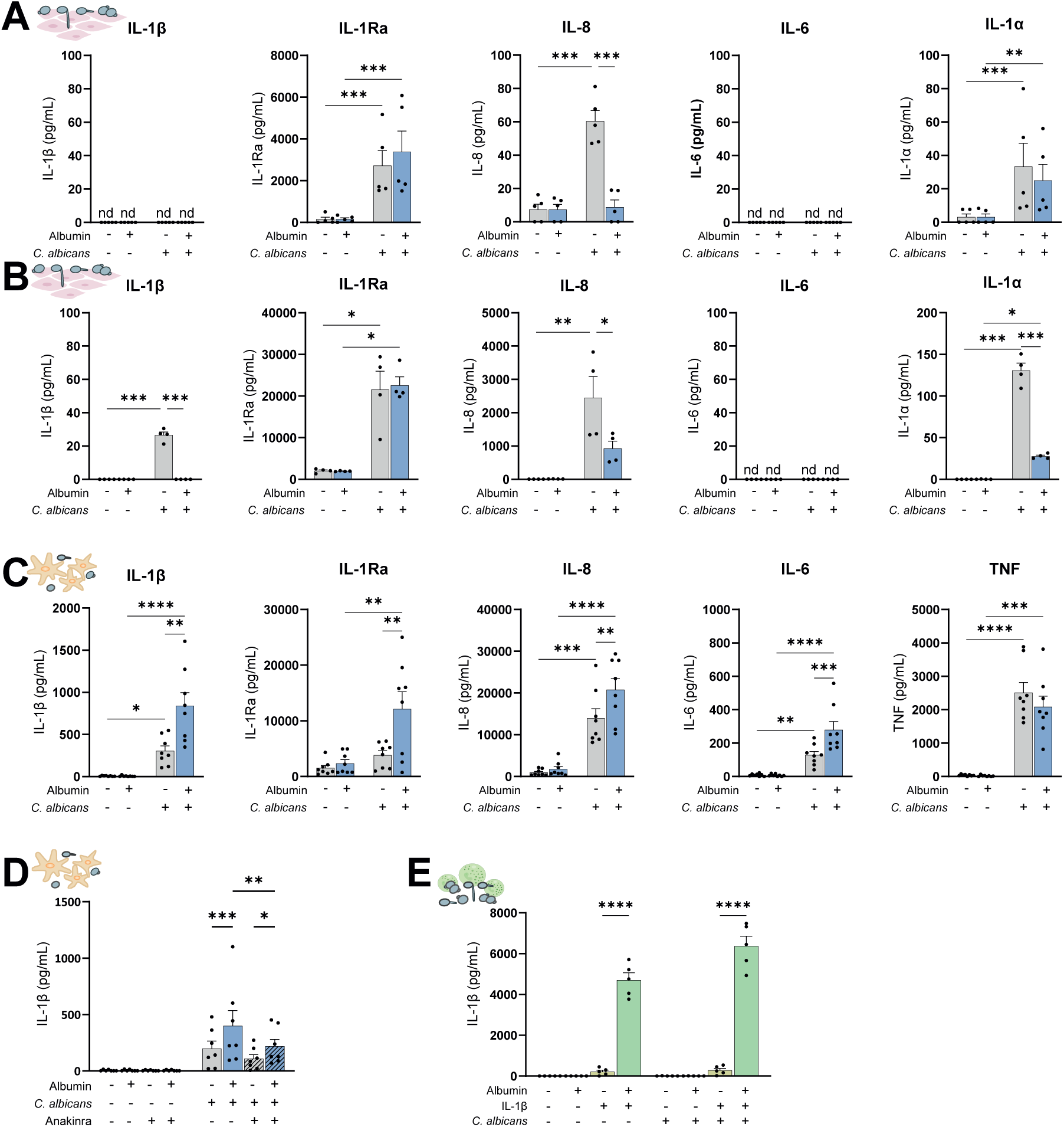
Albumin drives proinflammatory cytokine release by macrophages. **(A)** Cytokine release by A-431 cells (*n* = 5) in response to *C. albicans* infection in the presence or absence of albumin at 24 hpi. **(B)** Cytokine release by SW954 cells (*n* = 4) in response to *C. albicans* infection in the presence or absence of albumin at 24hpi. **(C)** Cytokine release by human monocyte-derived macrophages (*n* = 8) in response to *C. albicans* in the presence or absence of albumin. (**D**) IL-1β release by human monocyte-derived macrophages (*n* = 8) in response to *C. albicans* in the presence or absence of albumin and/or anakinra (*n* = 7). (E) IL-1β levels in culture supernatants of neutrophils (*n* = 5) in response to *C. albicans* in the presence or absence of albumin and/or supplemented IL-1β. Data are from at least three independently conducted experiments. Bars represent the mean + SEM with dots as individual biological replicates. Statistical analysis was done using a two-way ANOVA. * = *P ≤* 0.05, ** = *P ≤* 0.01, *** = *P ≤* 0.001, **** = *P ≤* 0.0001.

We used another cell line (SW954 cells) representing the vulvar epithelium. While these cells mounted modest IL-1α and IL-1β responses to *C. albicans* infection, the presence of albumin neutralized their release (**Figure 2B**). Albumin similarly suppressed IL-8 responses of SW954 cells. IL-6 remained undetectable in all conditions and IL-1Ra release was not changed by albumin.

In addition to epithelial cells, macrophages are important tissue-resident sentinels patrolling mucosal surfaces, including the vaginal mucosa (17–20). Therefore, we assessed the impact of albumin on macrophage responses to *C. albicans*. Human monocyte-derived macrophages (hMDMs) showed significantly increased cytokine responses to the combination of *C. albicans* and albumin (**Figure 2C**). The enhanced IL-1β, IL-1Ra, IL-8 and IL-6 release by hMDMs aligns with the correlations observed in RVVC patients (**Figure 1B**). IL-1β is notorious for amplifying its own release, therefore we sought to dampen this using recombinant IL-1Ra (Anakinra). Macrophage stimulation in the presence of albumin and anakinra returned the IL-1β levels back to similar levels as when albumin was absent (**Figure 2D**). Nevertheless, even when the self-amplifying loop is interrupted by Anakinra, IL-1β release was significantly boosted by albumin.

Following their recruitment to the vaginal mucosa, neutrophils will interact simultaneously with both albumin and *C. albicans*. Unlike macrophages, neutrophils failed to mount any detectable IL-1β responses to *C. albicans*, which was also not increased by the presence of albumin (**Figure 2E**). In line with previous observations, when stimulated with exogenous IL-1β, neutrophils fully degraded the cytokine to undetectable levels (35). Strikingly, the presence of albumin could prevent a substantial degradation (**Figure 2E**). Collectively, albumin-enhanced macrophage inflammatory responses to *C. albicans*, particularly IL-1β responses. Consequently, we further dissected the underlying mechanisms of how albumin exacerbates IL-1-driven inflammatory responses using macrophages.

### Albumin enhances *C. albicans*-induced NLRP3 inflammasome activation

Given the central role of IL-1β in driving innate and adaptive immune responses (36, 37), its processing and release is tightly regulated (37). In macrophages, IL-1β processing is a two-step process requiring priming and activation of the NLRP3 inflammasome (21) (**Figure 3A**). We hypothesized albumin increased *C. albicans*-induced NLRP3 inflammasome activation in macrophages, underlying their hyperinflammatory response (**Figure 2A**). Macrophages were primed with LPS to increase pro-IL-1β and pro-IL-18 levels, and subsequently stimulated with *C. albicans* to activate the inflammasome. Both IL-1β and IL-18 release induced by *C. albicans* increased by albumin (**Figure 3B**). Similarly, enhanced IL-1β release was observed in murine bone-marrow derived macrophages (BMDMs) stimulated with *C. albicans* in the presence of either human (**Figure 3C**) or murine albumin (**Supplementary Figure S4**), although less potently.

**Figure 3:**
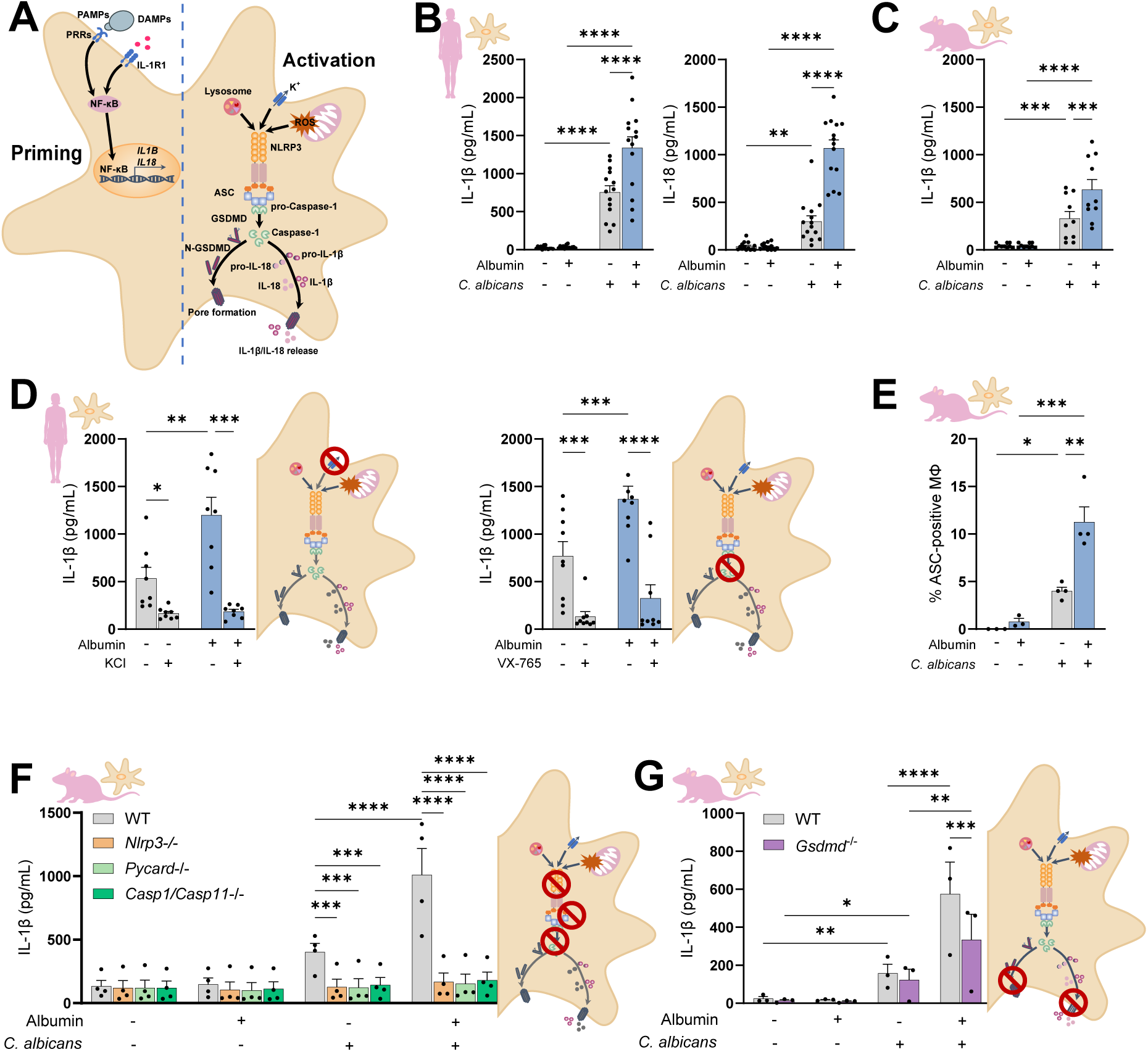
Albumin boosts *C. albicans-*induced NLRP3 inflammasome activation in macrophages. **(A)** Overview of NLRP3 inflammasome priming and activation. DAMPs = damage-associated molecular patterns. TLR = Toll-like receptor. PAMPs = pathogen-associated molecular pattern. **(B)** IL-1β and IL-18 release by hMDMs (*n* = 14 donors). **(C)** IL-1β release by mBMDMs (*n* = 10). **(D)** IL-1β release by hMDMs in the presence of the inflammasome inhibitors KCl (*n* = 8 donors) or VX-765 (*n* = 9 donors). **(E)** Percentage ASC-positive mBMDMs (*n* = 3 uninfected*, n* = 4 infected). **(F, G)** IL-1β release of mBMDMs deficient in the NLRP3 receptor (*Nlrp3*^−/−^), the adapter proteins ASC (*Pycard*^−/−^) or the proteolytic enzyme caspase 1 (*Casp1*^−/−^/*Casp11*^−/−^) (*n* = 4) or deficient in gasdermin D (*Gsdmd^−/−^*) (G). Bars represent the mean + SEM with dots as individual biological replicates: Individual donors (B, D) or mice (C, E - G) from at least three independently conducted experiments. Statistical significance was determined using a two-way ANOVA including with Holm-Šídák post-hoc test. * = *P ≤* 0.05, ** = *P ≤* 0.01, *** = *P ≤* 0.001, **** = *P ≤* 0.0001

Pro-IL-1β processing can either be caused by canonical NLRP3 inflammasome activation or non-canonical IL-1β processing (38). The NLRP3 inflammasome complex consists of three units: the sensor receptor domain NLRP3, the adapter protein ASC and the proteolytic enzyme caspase-1 (21) (**Figure 3A**). To determine whether albumin promotes activation of the canonical NLRP3 inflammasome, potassium efflux, an important trigger of NLRP3 activation, was inhibited by increasing the extracellular potassium concentration. Additionally, the role of caspase-1 was determined using the caspase-1 inhibitor VX-765. The albumin-enhanced IL-1β release by hMDMs was diminished by both KCl and VX-765 (**Figure 3D**).

Using BMDMs isolated from mice expressing the fluorescent citrine coupled to the ASC adapter protein, inflammasome assembly could be visualized (39), and a higher abundance of “ASC specks” was observed in the presence of albumin compared to the control condition (**Figure 3E**). BMDMs from mice deficient for each of the inflammasome components, the sensor Nlrp3 (*Nlrp3*^−/−^), the adapter protein ASC (*Pycard*^−/−^), and the proteases caspase-1 and −11 (*Casp1*^−/−^/*Casp11^−/^*^−^), failed to induce IL-1β release in the presence and absence of albumin (**Figure 3F**).

Besides pro-IL-1β processing, caspase-1-mediated gasdermin D (GSDMD) cleavage enables the formation of GSDMD pores within the macrophage membrane that facilitate IL-1β release as well as a cell death process called pyroptosis (40, 41). When stimulating gasdermin D (*Gsdmd^−/−^*) deficient murine BMDMs with *C. albicans*, the IL-1β release still increased in the presence of albumin, yet this increase was lower than in BMDMs from wild type (WT) mice (**Figure 3G**). This indicates that the NLRP3 inflammasome-mediated IL-1β release in the presence of albumin and *C. albicans* is partially, but not fully, GSDMD-dependent.

### Albumin-enhanced IL-1β release is driven by *C. albicans*

To ensure that the albumin-potentiated NLRP3 inflammasome activation is not exclusive to the *C. albicans* reference strain SC5314, the IL-1β response of macrophages to different *C. albicans* strains in the presence of albumin was quantified (**Figure 4A**).

**Figure 4:**
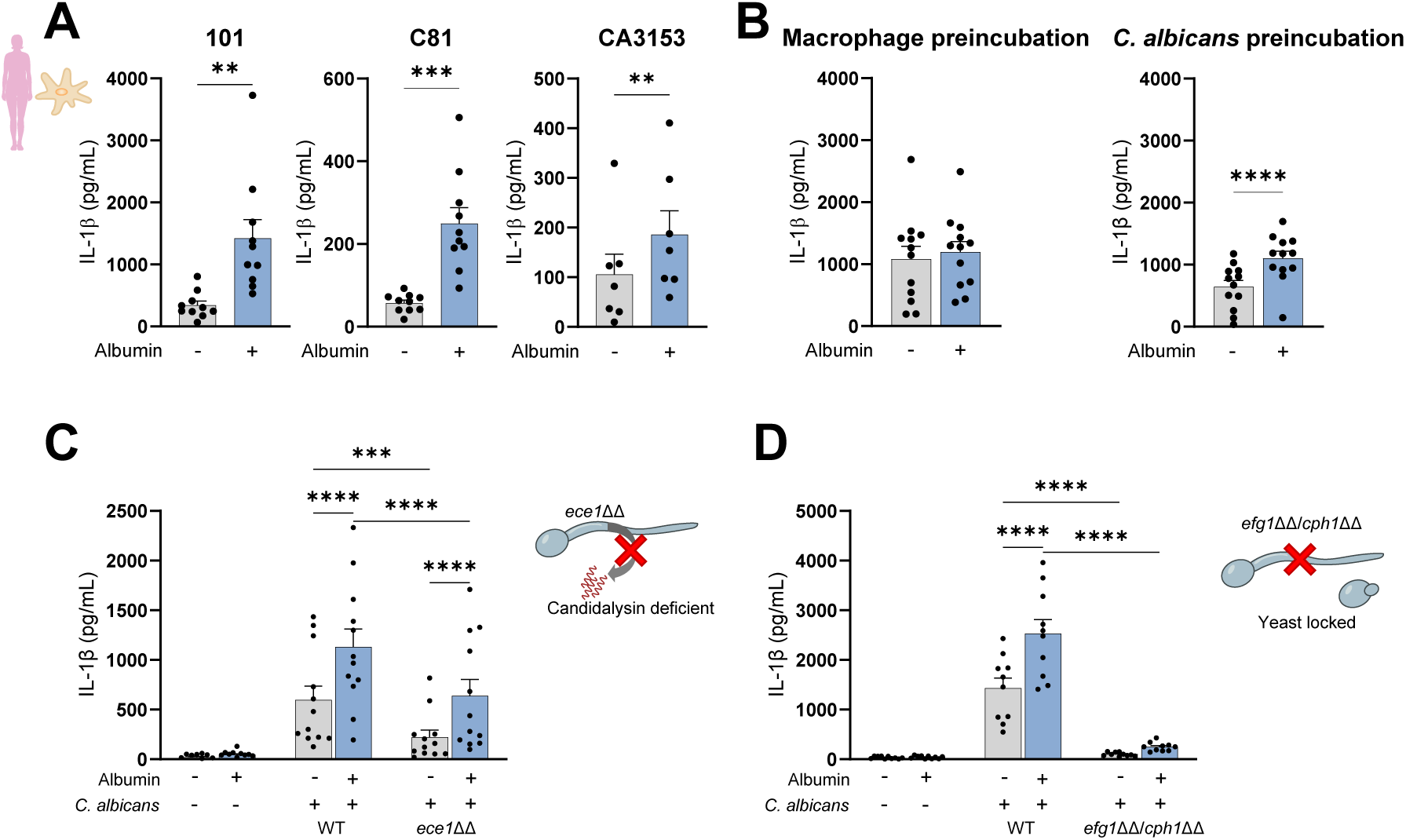
Albumin-enhanced IL-1β release relies on *C. albicans* and is independent of candidalysin. **(A)** IL-1β release by human macrophages in the presence or absence of albumin induced by the oral commensal isolate 101 (*n* = 10) and the vaginal isolates C81 (*n* = 10), CA3153 (*n* = 7). **(B)** IL-1β release of hMDMs in response to *C. albicans,* when either only hMDMs (left panel; *n* = 12) or *C. albicans* cells (right panel; *n* = 12) were preincubated with albumin prior to infection. **(C, D)** IL-1β release by hMDMs induced by *C. albicans* WT and a candidalysin-deficient *ece1*ΔΔ mutant (C; *n* = 12) or a yeast-locked *efg1*ΔΔ *cph1*ΔΔ mutant (D; *n* = 10). Bars represent the mean + SEM with dots as individual biological replicates of individual donors from at least three independently performed experiments. Statistical significance was determined using a paired t-test (A + B) or a two-way ANOVA with Holm-Šídák post-hoc test (C, D) * = *P ≤* 0.05, ** = *P ≤* 0.01, *** = *P ≤* 0.001, **** = *P ≤* 0.0001

By exclusively exposing either the macrophages or the fungus to albumin before infection, we found that albumin does not change the capacity of macrophage to activate the inflammasome (**Figure 4B, left panel**), but instead potentiated *C. albicans* to activate the inflammasome (**Figure 4B**, **right panel**).

To unravel how albumin causes *C. albicans* to induce stronger inflammasome activation, the role of selected virulence factors known to activate the NLRP3 inflammasome was investigated (42). Candidalysin, the major damage-inducing factor of *C. albicans*, induces inflammasome activation in macrophages (15). However, a strain lacking the candidalysin-encoding gene *ECE1* (*ece1*ΔΔ) still increased IL-1β release in the presence of albumin (**Figure 4C**). A yeast-locked *C. albicans efg1*ΔΔ*cph1*ΔΔ mutant poorly induced IL-1β release by hMDMs, whereas the cytokine release was not significantly increased by albumin (**Figure 4D**). Consequently, the albumin-enhanced IL-1β release by macrophages in response to *C. albicans* largely depends on the exposure to hyphae and hyphae-associated virulence factors.

### Albumin-enhanced IL-1β release relies on *C. albicans* secreted aspartic proteinase 1

*C. albicans* harbors a family of secreted aspartic proteinases (Saps) (13), of which some members have been described to mediate inflammasome activation (14, 43, 44). Individual *C. albicans* Sap deletion mutants for the yeast-associated Saps 1-3 and the hyphae-associated Saps 4-6 were assessed. Specifically, a *C. albicans sap1*ΔΔ mutant failed to increase IL-1β release in the presence of albumin (**Figure 5A**). This observation was reproduced using an independently generated *C. albicans sap1*ΔΔ mutant and was restored in a *sap1*ΔΔ + *SAP1* complemented strain (**Supplementary Figure S5**).

**Figure 5:**
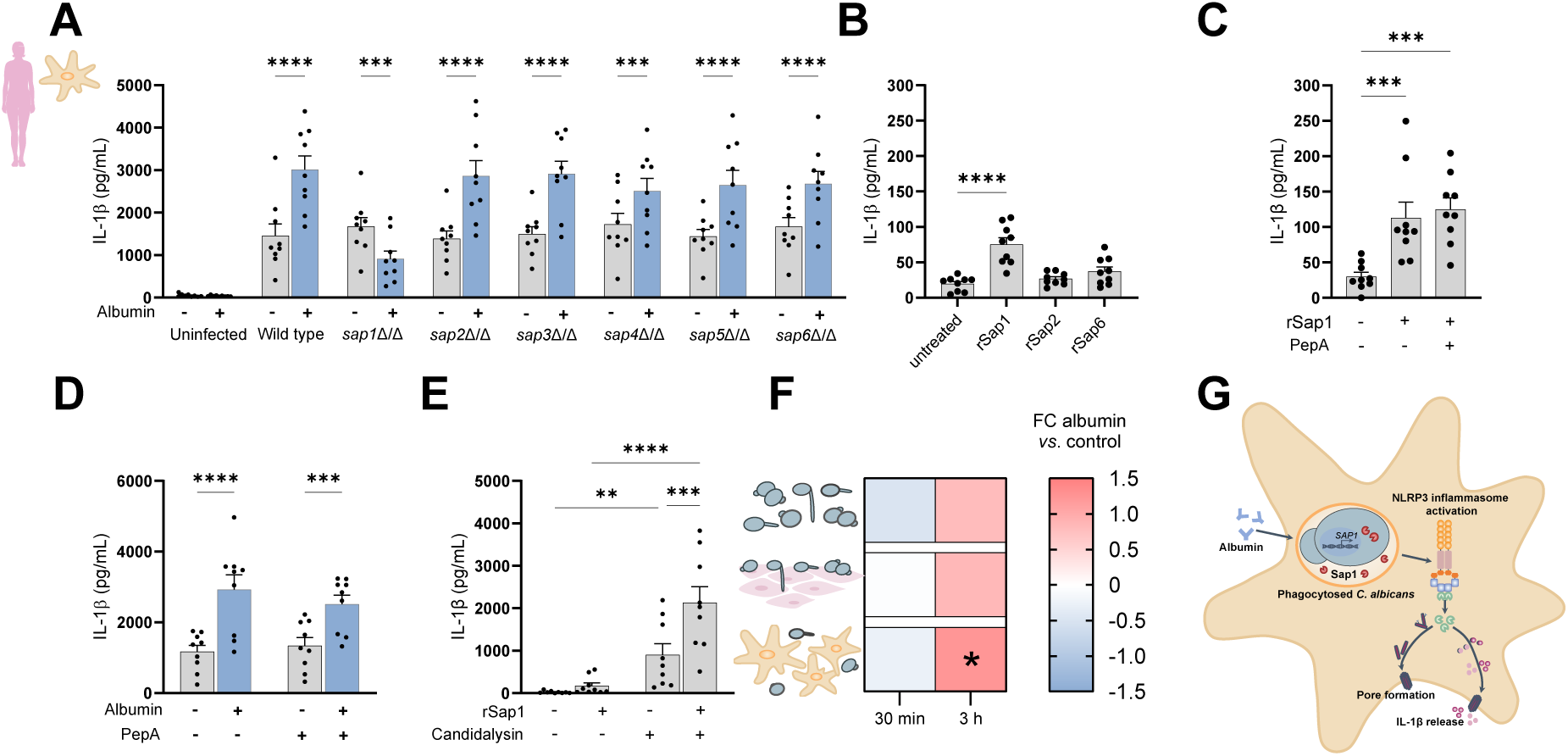
*C. albicans* Sap1 drives albumin-enhanced IL-1β responses. **(A)** IL-1β release by hMDMs induced by *C. albicans* WT or Sap1-6 deletion mutants (*n* = 9). **(B)** IL-1β release by hMDMs induced by recombinant Saps (*n* = 9). **(C + D)** IL-1β release by hMDMs induced by rSap1 (*n* = 9; C) or *C. albicans* (*n* = 9; D) in the presence or absence of pepstatin A (PepA). **(E)** IL-1β release by hMDMs induced by rSap1, synthetic candidalysin or the two combined (*n* = 9). **(F)** Normalized *SAP1* expression of *C. albicans* incubated in medium alone (*n* = 6), on VECs (*n* = 6) or with hMDMs (*n* = 6). Data were normalized to the condition without albumin and are shown as log_2_ fold change (FC). **(G)** Schematic showing albumin potentiated NLRP3 inflammasome activation by *C. albicans* Sap1. Data are from at least three independently conducted experiments. Bars represent the mean + SEM with dots as individual biological replicates. Statistical significance was determined using a one-way ANOVA including with Holm-Šídák post-hoc test (A - E) or a paired t-test (F). * = *P ≤* 0.05, ** = *P ≤* 0.01, *** = *P ≤* 0.001, **** = *P ≤* 0.0001.

To verify Sap1 as a NLRP3 inflammasome inducer in macrophages, hMDMs were stimulated with the recombinant Sap (rSap) 1 protein (45). In contrast to rSap2 and rSap6, rSap1 induced significant IL-1β release in LPS-primed hMDMs (**Figure 5B**). Consistent with a previous study (45), NLRP3 inflammasome activation was independent of the proteolytic activity of rSap1 as pepstatin A, a well-characterized Sap inhibitor, did not diminish IL-1β release (**Figure 5C**). In line with this, the albumin-enhanced IL-1β response to *C. albicans* was still evident in presence of pepstatin A (**Figure 5D**), suggesting that overall Sap proteolytic activity is not essentially required for the enhanced IL-1β release.

As candidalysin seemed to partially contribute to the elevated IL-1β responses to *C. albicans* in the presence of albumin (**Figure 4C**), we investigated how candidalysin and Sap1 act together on NLRP3 inflammasome activation. Indeed, the combination of rSap1 with synthetic candidalysin induced a synergistic IL-1β response (**Figure 5E**). As a short exposure of *C. albicans* to albumin prior to the macrophage encounter was sufficient to enhance its ability to induce IL-1β responses (**Figure 4B**), we hypothesized that albumin may induce *SAP1* mRNA expression. A tendency for increased *SAP1* expression in the presence of albumin was found after 3 h when *C. albicans* was grown alone or while infecting vulvovaginal epithelial cells (**Figure 5F**, **Supplementary Figure S6 A+B**). During the interaction with hMDMs, *SAP1* expression significantly increased in the presence of albumin (**Figure 5F**, **Supplementary Figure S6C**). Collectively, our results suggest that *C. albicans* increases *SAP1* expression in response to albumin, which activates the NLRP3 inflammasome in macrophages and promotes IL-1β responses (**Figure 5G**).

### Albumin promotes fungal fitness during confrontation with macrophages

As albumin acted on the fungus to elicit an elevated inflammatory response by macrophages, we systematically assessed how albumin altered *C. albicans-*macrophage interactions.

*C. albicans* (both WT and a *sap1*ΔΔ mutant), showed significantly improved survival rates when encountering macrophages in the presence of albumin. In contrast, a yeast-locked *C. albicans efg1*ΔΔ*cph1*ΔΔ mutant did not benefit from the presence of albumin (**Figure 6A**). We went through different stages of the interaction to pinpoint how albumin supports fungal survival within macrophages. Lower numbers of *C. albicans* cells were phagocytosed in the presence of albumin after 30 min, irrespective of the strain used (**Figure 6B**). Nevertheless, this difference was restored 2.5 h after infection with the *C. albicans* WT and *sap1*ΔΔ mutant, suggesting that albumin delays phagocytosis only at early stages (**Supplementary Figure 7 A + B**). Interestingly, albumin promoted the phagocytosis of the *efg1*ΔΔ*cph1*ΔΔ mutant 2.5 h after infection (**Supplementary Figure S7C**).

**Figure 6:**
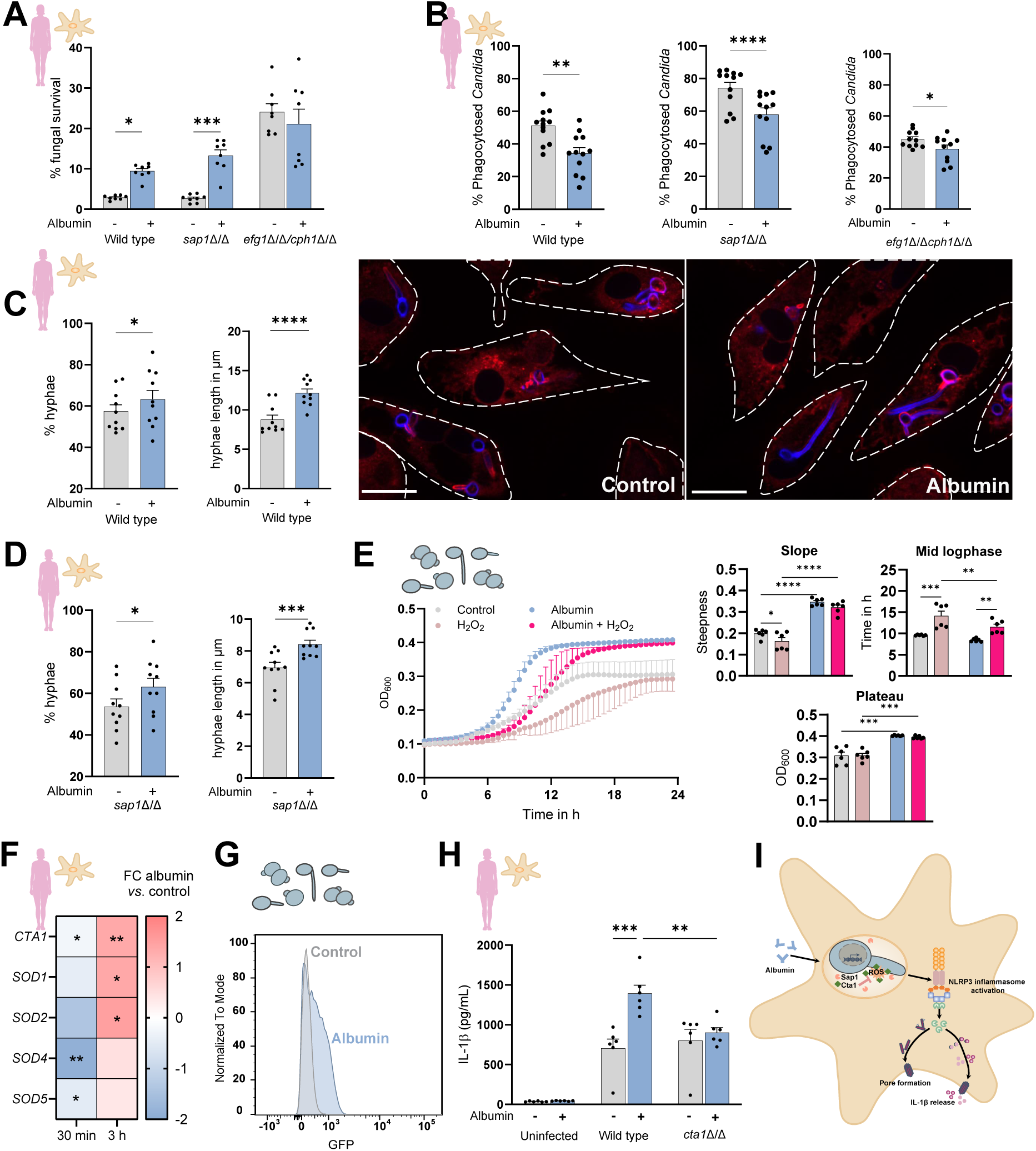
Albumin promotes fungal fitness within macrophages. **(A)** Fungal survival 3 h after infection of hMDMs (*n* = 12). **(B)** *C. albicans* phagocytosis by hMDMs 30 min after infection (*n* = 12). **(C + D)** Percentage and length of *C. albicans* hyphae 2.5 h after macrophage infection WT (C; *n* = 10 donors) or *sap1*ΔΔ (D; *n* = 10). Representative images with scale bars indicate 20 μm. **(E)** *C. albicans* growth under oxidative stress (*n* = 6). **(F)** Normalized expression of *C. albicans* oxidative stress resistance genes during the interaction with hMDMs after 30 min (*n* = 10) and 3 h (*n* = 8). Expression data is shown as log_2_ fold change (FC) of albumin compared to the control without albumin. **(G)** Representative histograms (normalized to mode) of *p-cta1-GFP* mean fluorescence intensity. **(H)** IL-1β release by hMDMs (*n* = 6) induced by *C. albicans* WT and *cta1*ΔΔ mutant. **(I)** Schematic showing albumin-promoted survival of *C. albicans* within macrophages allowing the expression of Sap1. Bars represent the mean + SEM with dots as individual biological replicates. Data are from at least three independently conducted experiments. Statistical significance was determined using paired t-tests (A-D, F) or by one-way ANOVA with Holm-Šídák post-hoc test (E, H). * = *P ≤* 0.05, ** = *P ≤* 0.01, *** = *P ≤* 0.001, **** = *P ≤* 0.0001.

Quantification of the fungal germination rate and hyphae length within macrophages suggested that albumin enables stronger filamentous growth (**Figure 6 C+D**). Intriguingly, in the absence of macrophages, *C. albicans* formed shorter hyphae in response to albumin (**Supplementary Figure S8**), suggesting that albumin specifically promotes filamentation within macrophages. This hints at an interference with the macrophage capacity to control *C. albicans* filamentation within the phagosome.

Albumin is known for its antioxidative properties (46), which could impair the oxidative environment of the phagolysosome, a crucial process in killing *C. albicans*. Accordingly, the growth-inhibiting effect of H_2_O_2_, indicated by a delayed time to mid log-growth, was restored by albumin (**Figure 6E**). Several oxidative stress genes were significantly downregulated 30 min after exposing the fungus to macrophages in the presence of albumin but upregulated after 3 h (**Figure 6F**, **Supplementary Figure S9A**). In line with this, using a reporter strain in which GFP expression is controlled by the promotor of the *C. albicans* catalase 1 gene (*CTA1*) (47), we showed that the presence of albumin activates *CTA1* promotor activity (**Figure 6G**, **Supplementary Figure S9 B+C**). Interestingly, a *C. albicans* mutant lacking the catalase 1 gene (*cta1*ΔΔ), a crucial factor for detoxifying hydrogen peroxide, lost its ability to enhance IL-1β release in the presence of albumin (**Figure 6H**).

Collectively, albumin promotes *C. albicans* survival within macrophages through supporting its oxidative stress resistance, leading to stronger intraphagosomal filamentation. This permits the increased expression of *SAP1* as an inflammasome inducer (**Figure 6I**).

### Albumin causes dysfunctional neutrophil activation and fungal clearance

Based on the albumin-enhanced macrophage responses to *C. albicans*, we hypothesized that this could translate to different downstream neutrophil activation. Primary human neutrophils stimulated with hMDM supernatants showed significantly increased surface expression of the multifunctional integrin CD11b and the degranulation marker CD66b. This indicated an elevated neutrophils activation, when macrophages upstream had been exposed to *C. albicans* and albumin (**Figure 7A**, gating strategy in **Supplementary Figure S10**).

**Figure 7:**
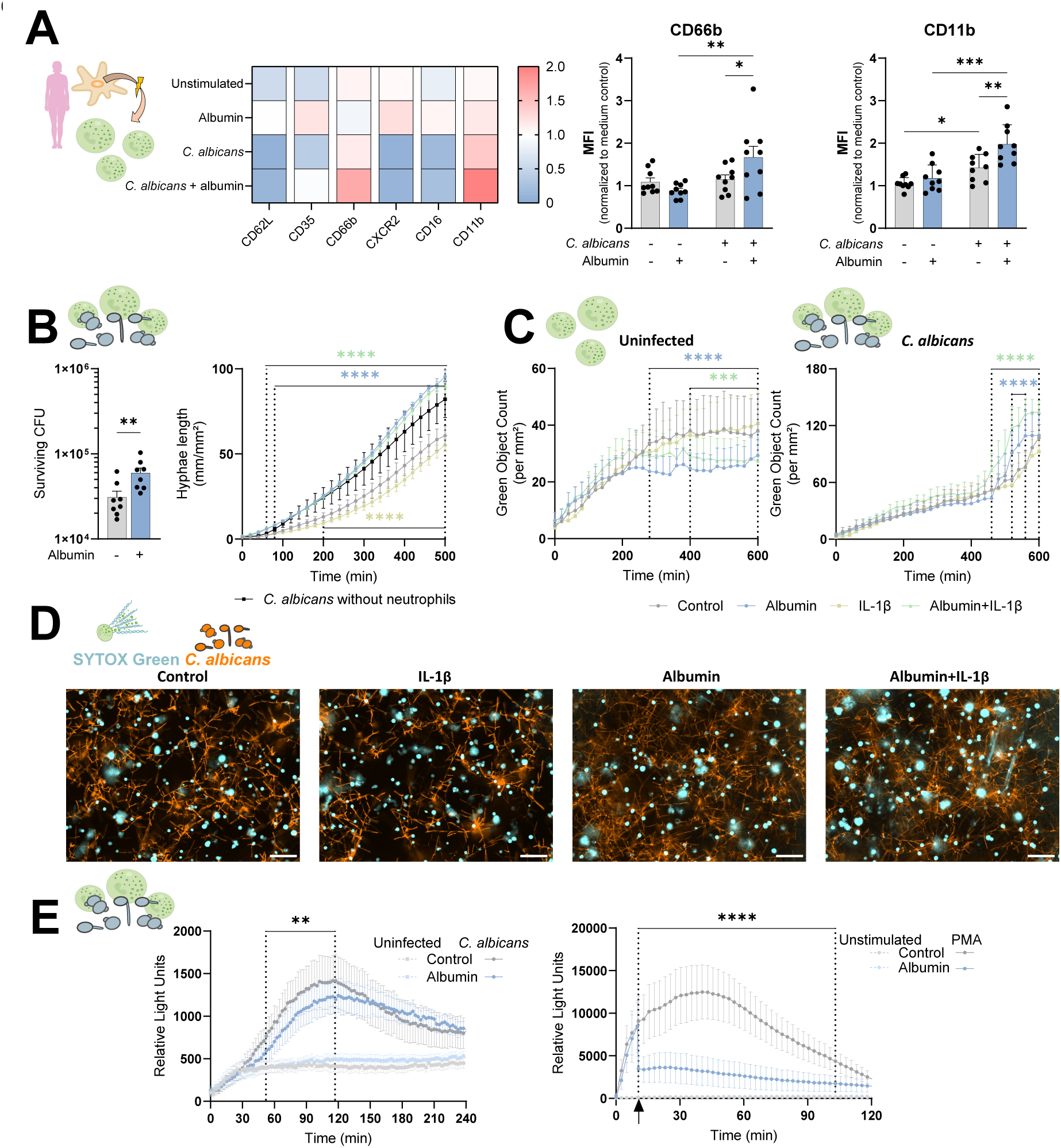
Albumin increases downstream neutrophil activation and compromises fungal clearance. (**A**) Neutrophil activation marker expression after stimulation with supernatants from macrophages challenged with *C. albicans* in the presence or absence of albumin (neutrophils *n* = 3 donors were stimulated with supernatants from *n* = 3 macrophage donors). Overview heatmap of all markers (left panel). Additionally, mean fluorescence intensity (MFI) of CD66b and CD11b is shown as bar graph with SEM (right panel). **(B)** Remaining fungal CFU following encounter with neutrophils in the presence or absence of albumin (left panel, *n* = 8, mean + SEM with individual biological replicates as dots). Fungal hyphal growth assessed by live-cell imaging in the presence or absence of neutrophils with or without albumin and supplemented IL-1β (Right panel, *n =* 5 donors). (**C**) Spontaneous (left) and *C. albicans*-induced (right) formation of neutrophil extracellular traps in the presence of albumin (5 mg/mL) and/or IL-1β (10 ng/mL) quantified by live-cell imaging. (**D**) Images of the live-cell analysis of fungal outgrowth and NETosis at 8 hpi. (**E)** Luminol conversion by reactive oxygen species produced by neutrophils in response to *C. albicans* in the presence or absence of albumin (left panel, *n* = 6) or in response to PMA with albumin injection after 10 min (right panel, *n* = 7). Statistical analysis was done using a 2-way ANOVA (A, B right panel, C, E) or paired t-test (B left panel). * = *P ≤* 0.05, ** = *P ≤* 0.01, *** = *P ≤* 0.001, **** = *P ≤* 0.0001.

As neutrophils are dysfunctional during RVVC (6, 8), we also investigated how albumin impacts their antifungal activity. Fungal clearance capacity of neutrophils improves by IL-1β stimulation (**Figure 7B**). Yet, albumin compromised the ability of neutrophils to restrict hyphal outgrowth and even overruled any IL-1β-derived benefits for antifungal activity (**Figure 7B, D**). Neutrophil extracellular trap formation (NETosis) is an important mechanism by which neutrophils attack fungal hyphae (48), yet this is dysregulated during VVC (49, 50). While albumin as well as the combination of albumin with IL-1β supported viability of uninfected neutrophils, both conditions promoted NETosis during infection (**Figure 7C, D**). The production of reactive oxygen species (ROS) is an important mechanism for fungal clearance by neutrophils. In line with the compromised neutrophil-mediated killing and attenuation of oxidative responses by albumin in macrophages, ROS activity derived from neutrophils was compromised by albumin (**Figure 7E**).

## DISCUSSION

Vulvovaginal candidiasis is characterized by dysfunctional hyperinflammatory responses (6, 8). Vaginal albumin concentrations increased upon *C. albicans* infection and correlated with the presence of proinflammatory cytokines in RVVC patients. We showed that this is not merely a correlation with inflammation, but also that albumin causally exacerbates proinflammatory responses of macrophages towards *C. albicans,* downstream resulting in increased neutrophil activation. Yet, both neutrophils and macrophages exhibit dysfunctional fungal clearance in the presence of albumin. Mechanistically, we found that albumin improves *C. albicans* fitness against macrophages and neutrophils by boosting fungal ROS resistance, which depends on albumin-induced oxidative resistance and albumin’s direct suppression of ROS. In the presence of albumin, *C. albicans* has increased expression of *SAP1*, which we identified as the factor enhancing NLRP3 inflammasome activation in macrophages. Collectively, we demonstrate a role for albumin in NLRP3-inflammasome-driven hyperinflammation and dysfunctional innate fungal clearance associated with VVC.

Macrophages seem an overlooked cell type in VVC, but are actually abundant in the vaginal mucosa (17–20). We established macrophages as plausible source of the VVC-driving cytokine IL-1β, as both epithelial cells of that niche and recruited neutrophils barely released any IL-1β compared to macrophages, when encountering *C. albicans*. This release was enhanced by the presence of albumin. With evidence on several layers, we show that albumin increases *C. albicans*-induced NLRP3 inflammasome activation. Albumin delayed *C. albicans* phagocytosis by macrophages, and compromised their capacity to control intracellular fungal survival. In line with the ability of albumin to neutralize reactive oxygen species (46), we observed that albumin increases the expression of oxidative stress resistance genes. Besides its effect on oxidative stress, albumin promotes global metabolic adaptations in *C. albicans* and can help it to overcome iron limitation (30, 51). Therefore, albumin may additionally serve as an iron source to overcome nutritional immunity the phagosome. Likely, a combination of these effects enables *C. albicans* to form longer filaments and to survive within macrophages.

*C. albicans*-induced IL-1β release by MDMs was dependent on fungal filamentation, as the yeast-locked *efg1*ΔΔ*cph1*ΔΔ mutant induced poor IL-1β responses, even in presence of albumin. After processing, bioactive IL-1β is released by gasdermin-D pore formation (a host-driven process) or macrophage lysis induced by fungal filamentation and candidalysin (15, 52, 53). This explains why a *C. albicans ece1*ΔΔ mutant generally induced lower IL-1β release in the presence of albumin. Similarly, BMDMs of *Gsdmd^−/−^* mice challenged with *C. albicans* released less IL-1β in the presence of albumin compared to WT BMDMs. Like previously suggested (52), this underscores redundancy between candidalysin and gasdermin-D mediated IL-1β release.

We pinpointed *C. albicans* protease Sap1 as the driver of elevated NLRP3 inflammasome activation. Albumin increased *SAP1* expression, particularly during the encounter with macrophages. Linking this to clinical VVC, *SAP1* was found to be upregulated in *C. albicans* isolates from VVC patients but not in isolates from healthy women (54). The general importance of Saps for *C. albicans* immunopathogenicity during VVC has been shown in numerous studies (7, 43, 54). Sap2, Sap5 and Sap6 are characterized inflammasome inducers (14, 22, 44), yet also Sap1 was shown to increase IL-1β release in monocytes (45). We observed that, stimulation of human macrophages with recombinant Sap1 induces higher IL-1β release than with Sap2 or Sap6. Our data complements previously published literature by underscoring the role of *C. albicans* secreted aspartic proteases in IL-1β responses during VVC.

The NLRP3 inflammasome drives neutrophil-mediated inflammation during (R)VVC (7, 22, 23). Such IL-1-driven inflammatory responses during *C. albicans* infections trigger neutrophil recruitment, which normally mediate resistance to infection (55, 56). However, during (R)VVC neutrophils are believed to aggravate disease than effectively clear the infection (4, 6, 8). We found that albumin compromises the ability of neutrophils to effectively clear *C. albicans*. This is associated with the antioxidant effects of albumin neutralizing neutrophil-derived ROS directly, as well as increasing fungal oxidative stress resistance. Importantly, the attenuated capacity to antagonize fungal growth was despite increased NET formation. This fits the observation of a high abundance of NETs in vaginal discharges of VVC patients (49). Mechanistically, albumin modified by neutrophil MPO-derived hypochlorous acid has been previously identified as a trigger of NETosis (57).

Our findings, in line with previous studies, reflect the complexity and multifactorial pathogenesis of VVC. The immunopathogenesis of VVC can hardly be pinpointed to a single factor but is a complex combination of different factors and effects on the host side as well as the fungal side (6, 8). By combining observations from RVVC patients and a murine VVC model with *in vitro* mechanistic dissection, we identified albumin as a catalyst of IL-1β-mediated inflammation that may underlie inflammation in VVC patients.

## Materials & Methods

### Ethics statement for primary human & murine macrophages and mouse in vivo experiments

All methods were conducted in accordance with relevant guidelines and regulations. Experimental protocols for murine *in vivo* studies were approved by the Ministry of Health (Authorization N. 725/2019-PR) and previously certified by the animal ethics committee ‘OPBA’ from the University of Perugia, Italy.

Human blood was obtained from healthy volunteers who gave written informed consent in compliance with the Declaration of Helsinki. The Jena institutional ethics committee approved the procedure (Ethik-Kommission des Universitätsklinikums Jena, Permission No. 2207–01/08). Murine bone marrow was obtained in compliance with the German animal protection law and the collection of bone marrow was approved by the local ethics committees of the University clinics of Bonn and Erlangen, respectively.

### Collection of human vaginal fluids

Two RVVC patient cohorts were used. The first cohort (RVVC_1_), consists of RVVC patients (*n* = 19) and control volunteers (*n* = 15) (32), who enrolled in a study approved by the University Ethics Committee of Perugia (Prot. 2012-028) and provided written informed consent for study participation. Vaginal fluid collection was performed at the S. Maria Della Misericordia Medical Center (Perugia, Italy). Cervicovaginal samples were obtained from all participants by instilling 3 mL of sterile saline into the posterior vagina, mixing the saline with secretions and withdrawing the solution with a syringe. All vaginal washes were centrifuged at 12 000 × *g* for 10 min to separate the mucus from the PBS wash solution shortly after collection and washes were immediately frozen at −20°C (32). The study population included Caucasian women diagnosed with at least three documented VVC episodes in a year, which were microbiologically validated, and caused by *C. albicans.* Controls consisted of age-matched healthy Caucasian women with no gynecological complaints, no history of vaginal candidiasis infection, and who were culture-negative for vaginal pathogens. Exclusion criteria were pregnancy, diabetes mellitus, endocrine or immune deficiency disorders, use of immunosuppressive medications, antibiotics or high estrogen content contraceptives, chemotherapy or prior hysterectomy (32).

The second cohort (RVVC_2_) served as a validation cohort. Vaginal lavages (*n* = 129) from a previous prospective multicentric follow-up study performed to evaluate the effectiveness of individualized decreasing fluconazole maintenance therapy to prevent relapses of *Candida* spp. vulvovaginitis were provided (ReCiDiF trial) (33). Vaginal rinsing with 2 mL of 0.9% sodium chloride was performed by flushing and re-aspirating the fluid through a 0.5-mm wide, 6-cm long needle in the left, central, and right upper vaginal vaults (34).

### Experimental design of murine VVC model

Female C57BL/6J mice were treated subcutaneously with 0.2 mg of β-estradiol 17–valerate (Merck) dissolved in 100 µL of sesame oil (Merck) 24 h before vaginal infection and on the day of vaginal infection. Estrogen administration continued once per week until the end of the experiment to maintain pseudoestrus. Mice were inoculated intravaginally with 20 µL of phosphate-buffered saline (PBS) or PBS containing 2 × 10^9^/mL viable *C. albicans* SC5314 yeast cells from early–stationary-phase cultures (i.e., 18 h of culture at 37 °C on Sabouraud-dextrose agar with chloramphenicol plates). Vaginal lavages were conducted with sterile PBS with repeated aspiration and agitation (total 100μL). The course of infection was monitored in individual mice by culturing 100 µL of serially diluted (1∶10) vaginal lavages on Sabouraud-dextrose agar with chloramphenicol. CFUs were enumerated after incubation at 37 °C for 24 h and expressed as log_10_ CFU/mL of lavage fluid.

### Murine histology

The reproductive tract was collected and fixed in 10% neutral buffered formalin, trimmed in transverse sections to include portions of the vagina, cervix, and uterus and embedded in paraffin. Histologic slides were prepared using routine methods and 4 μm-thick sections were stained with hematoxylin and eosin or Grocott’s methenamine silver stain for enhanced fungal visualization. The stained slides were observed on a BX53 system microscope (Olympus). Alternatively, immunofluorescence to visualize neutrophil tissue influx and fungal presence was performed. For this, 5 µm-thick sections were cut onto adhesive slides (Superfrost Plus Adhesion, Epredia) and deparaffinated by submersion in a row of xylene (2 × 10 min), 100% ethanol (2 × 1 min), 95% ethanol (1 min), 75% ethanol (1 min), tap and distilled water (1 min each). Samples were encircled with an ImmEdge hydrophobic barrier PAP pen (Vector Laboratories) and after brief drying, slides were washed with PBS including 0.3% Triton-X100. They were blocked with 5% goat serum in PBS with 0.3% Triton-X100 for 90 min and, without prior antigen-retrieval, overnight stained with 80 µl primary antibody mix including PBS, 1% BSA, 0.3% Triton-X100, 10 µg/ml anti-*Candida* (polyclonal rabbit, BP1006, Acris Antibodies/OriGene Technologies) and 5 µg/ml AlexaFluor-647-coupled anti-Ly6G (clone 1A8, #127610, Biolegend) at 4 °C inside a humid chamber. Samples were washed thrice and secondary stained with 5 µg/ml goat-anti-rabbit IgG-AlexaFluor 488 (A-11008, Invitrogen, Thermo Fisher Scientific) and 5 µg/ml DAPI for 90 min at RT in the humid chamber. Samples were washed thrice and embedded in Fluoroshield (Invitrogen). Images were acquired on an Axio Observer Z1 (Carl Zeiss Microscopy).

### Cytokine and albumin measurements

Human albumin levels were determined using human albumin ELISA kits (Abcam ab108788 for RVVC_1_ or ab108787 for RVVC_2_), according to manufacturer instructions. In samples of the RVVC_1_ cohort and the healthy control group, levels of IFN-α, IFN-β, IL-1α (IL-1F1), IL-1β (IL-1F2), IL-1Ra (IL-1F3), IL-6, IL-8 (CXCL8), IL-17A, GM-CSF and TNF were quantified using a human discovery Luminex© assay (R&D Systems). For the RVVC_2_ cohort and *in vitro* experiments with human macrophages or vaginal epithelial cells, cytokine levels (IL-1α, IL-1β, IL-1Ra, IL-6, IL-8, IL-17 and TNF) in vaginal lavages were quantified by DuoSet ELISA (R&D systems) (34).

Murine albumin levels were determined using a murine albumin ELISA kit (Abcam). Vaginal levels of murine IL-1β (IL-1F2) and MPO were quantified using DuoSet ELISAs (R&D Systems).

### Strains and culture conditions

*C. albicans* strain SC5314 was used as WT strain if not indicated otherwise. BWP17 + CIP30 was used the reference strain for experiments with the candidalysin-deficient strain (*ece1*ΔΔ) (15). Other *C. albicans* strains used in this study included the oral commensal isolate 101 (kindly provided by Salomé LeibundGut-Landman, University of Zurich) (58), the vaginal isolate C81 (kindly provided by Julian Naglik, King’s College London), the vaginal isolate CA3153 (kindly provided by Patrick van Dijck, KU Leuven), a yeast-locked *C. albicans* mutant (*efg1*ΔΔ /*cph1*ΔΔ) (59) and *C. albicans SAP1*-*SAP6* deletion strains (*sap1*ΔΔ, *sap2*ΔΔ, *sap3*ΔΔ, *sap4*ΔΔ, *sap5*ΔΔ, *sap6*ΔΔ) (60). CAI4 + Clp10 (61) was used as reference strain for the validation experiment with an independently generated *SAP1* deletion strain (*sap1*ΔΔ) (62) and *SAP1* complemented strain (*sap1*ΔΔ + SAP1) (62) as well as for the *CTA1* deficient strain (*cta1ΔΔ*) (63). Further, *C. albicans* GFP reporter strains to quantify the promotor activity of *ACT1* (pACT1-GFP) (61), *CTA1* (pCTA1-GFP) (64) and *SOD5* (pSOD5-GFP) (61) were used. Strains were streaked on Yeast Peptone Dextrose (YPD) agar plates and maintained for two weeks. Single colonies were inoculated in liquid YPD medium and grown overnight shaking at 30 °C and 180 rpm. Fungal cells were washed three times with phosphate-buffered saline (PBS, pH 7.4) and adjusted to the required cell concentration.

### Albumin solutions

Human albumin was obtained from Sanquin Plasma Products B.V. (Albuman) and murine albumin was used from Innovative Research. If not indicated differently, human albumin was used for experiments. Murine albumin was used for selected experiments. Albumin was used at a final concentration of either 5 mg/mL or 10 mg/mL, exact concentrations used are given in the description of the respective assay.

### In vitro vaginal epithelial infection model

To mimic vulvovaginal mucosal infection, A-431 cells (VECs, Deutsche Sammlung von Mikroorganismen und Zellkulturen DSMZ no. ACC 91) were used. The cell line has been authenticated *via* commercial STR profiling (Eurofins Genomic) and checked for mycoplasma contaminations using a PCR mycoplasma test kit (PromoKine). VECs were maintained in RPMI-1640 medium containing L-glutamine (Thermo Fischer Scientific) and supplemented with 10% heat-inactivated fetal bovine serum (FBS; Bio & SELL) at 37 °C and 5% CO_2_. For infection experiments, 1 × 10^5^ VECs per mL were seeded in 96-well plates (200 µL/well) or 6-well plates (3 mL/well) and cultured for 2 days (37 °C, 5% CO_2_). At the day of infection, VECs were infected with *C. albicans* strains at a multiplicity of infection (MOI) of 1, unless stated otherwise, in RPMI-1640 without FBS and with or without 5 mg/mL albumin. Infected VECs were incubated for 24 h (37 °C, 5% CO_2_). Afterwards, samples were centrifuged at 200 × *g* for 10 min, supernatants were collected and stored at −20 °C until cytokine quantification.

### Human macrophage differentiation

Human monocytes were isolated and differentiated to macrophages as described previously (15). In brief, human peripheral blood mononuclear cells (PBMCs) were isolated from buffy coats using density centrifugation with lymphocyte separation medium (1.077 g/mL; Capricorn scientific). After washing the PBMCs twice in PBS, magnetic automated cell sorting (autoMACS, MiltenyBiotec) was used for monocyte isolation using anti-CD14-labeled beads. Human monocyte-derived macrophages (hMDMs) were differentiated by seeding 1 × 10^7^ monocytes in 175 cm^2^ cell culture flask in RPMI-1640 medium containing L-glutamine and supplemented with 10% heat-inactivated FBS and 50 ng/mL recombinant human M-CSF (ImmunoTools) for 7 days at 37 °C and 5% CO_2_ with a medium refreshment after 5 days. Macrophages were detached by adding PBS containing 10 mM EDTA, incubating the cells for 30 min (37 °C, 5% CO_2_) and subsequent cell scraping. If not stated differently, hMDMs were seeded at 4 × 10^4^ cells/well in 96-well plates. For the collection of RNA, 6 × 10^5^ cells/well were seeded in 6-well plates. Macrophages were rested over night before infection.

### Murine macrophage cultures

Murine bone-marrow-derived macrophages (BMDMs) were generated from bone marrow cells flushed from femurs and tibiae of SPF C57BL/6J WT, *Nlrp3*^−/−^(65), *Pycard*^−/−^ (66), *Casp1*^−/−^/ *Casp11^−/−^* (67, 68) or *Gsdmd*^−/−^ (40) deficient or R26-CAG-ASC-citrine reporter mice (39) (JAX stock #030744). Isolated cells from one leg were seeded into a 175 cm^2^ cell culture flask. Differentiation was done for 7 days at 37 °C and 5% CO_2_ in RPMI-1640 medium containing 10% FBS, 40 ng/mL recombinant murine M-CSF (ImmunoTools) and 1 × Penicillin/Streptomycin (100 U/mL; Gibco) with a medium exchange on day 5. Detachment and seeding of murine BMDMs was done as described for human MDMs.

### Macrophage stimulations

For cytokine assays, human MDMs were stimulated with *C. albicans* at MOI 1. Albumin was added at concentration of 10 mg/mL. Samples were incubated at 37 °C and 5% CO_2_. After 24 h, samples were centrifuged at 200 × *g* for 10 min, supernatants were collected and stored at −20 °C until cytokine quantification.

Inflammasome activation was assessed with human MDMs or murine BMDMs as previously published (15). Macrophages were primed with 50 ng/mL lipopolysaccharide (LPS; from *Escherichia coli*, Sigma) 2 h before infection if not mentioned differently. For inhibitor experiments, Anakinra (recombinant human IL-1Ra, 10 μg/mL, Kineret), potassium chloride (25 mM; Merck) or the caspase-1 inhibitor VX-765 (50 μg/mL; Invivogen) were added 1 h prior to infection. Subsequently, 10 mg/mL albumin was added and macrophages were infected with *C. albicans* at MOI 10. For preincubation experiments, macrophages or *C. albicans* were exposed to 5 mg/mL human albumin 1 h prior to infection. For *C. albicans* preincubation, fungal cells were harvested by centrifugation at 3,000 × *g* washed twice with PBS and adjusted to the concentration for infection. For macrophage preincubation, supernatant containing albumin was removed and cells were washed once with RPMI-1640 before infection. During indicated experiments, proteinase activity was inhibited using the proteinase inhibitor pepstatin A (Sigma; 15 μM). Infected macrophages were incubated for 5 h (37 °C, 5% CO_2_) before harvesting the supernatants as described above. For assays with recombinant Saps, Sap1, Sap2, or Sap6 were added at a concentration of 20 μg/mL after LPS priming and assay time was extended to 24 h. For combination experiments of Sap1 with candidalysin, 20 μM synthetic candidalysin (Peptide Synthetics) were used.

### Neutrophils isolation

Neutrophils were isolated from freshly drawn venous blood as described previously (35). Blood was diluted 1:1 with sterile PBS, and cells were separated by layering lymphocyte separation medium (1.077 g/mL; Capricorn scientific) under the blood and centrifugation for 30 min at 700 × *g* and room temperature. Plasma, PBMCs, and lymphocyte separation medium were removed and hypotonic lysis buffer was added to the layer of erythrocytes and granulocytes. Samples were incubated on ice for 15 min and subsequently centrifuged at 300 × *g* and 4 °C for 10 min. Supernatant was removed and the lysis step was repeated for 10 min. The pellet was washed twice with ice-cold PBS and centrifuged at 300 × *g* and 4 °C for 10 min. After washing, cells were resuspended in RPMI-1640 and adjusted to the required assay concentration.

### Flow cytometric analysis of neutrophil activation

Neutrophil activation assay was done as described previously with minor modifications (9). Human MDMs were stimulated with RPMI-1640 with or without 10 mg/mL albumin and infected with *C. albicans* infection at a MOI of 1. After 24 h, macrophage supernatants were used to stimulate human neutrophils. Neutrophils were seeded at a density of 2 × 10^5^ cells/well in a 96-well plate. After 2 h of stimulation, neutrophils were washed using cold FACS buffer (PBS containing 2% FBS). Neutrophils were preincubated with Fc-Block Human TruStain FcX (BioLegend) and stained using fluorophore-linked antibodies against CD15-APC/Fire750 (SSEA-1) and the activation markers CD62L-AlexaFluor647 (DREG-56), CD35-FITC (E11), CD66b-PE (G10F5), CXCR2/CD182-PE-Cy7 (5E8) CD16-PerCP-Cy5.5 (3G8) and CD11b-BV421 (ICR44; all from Biolegend). Dead cells were stained using the fixable viability dye eFluor506 (Invitrogen, Thermo Fisher Scientific). Neutrophil were stained for 20 min at 8 °C in the dark, cells were washed in FACS buffer and filtered through a 70 μm mesh before measurement on a FACSVerse Cell Analyzer (BD Biosciences). FlowJo v.10. was used for analysis, gating strategy is depicted in **Supplementary Figure S9**.

### Fungal killing by neutrophils

Human neutrophils were seeded in 96-well plates at a concentration of 5 × 10^4^ cells/well and infected with *C. albicans* MOI 0.5 in the presence or absence of 5 mg/mL albumin. Cells were incubated at 37 °C and 5% CO_2_. After 3 h, the content of the wells was scraped, resuspended and transferred to a collection tube. Afterwards, sterile H_2_O was added to each well and remaining content of the wells was lysed, resuspended and transferred to the respective collection tube. The procedure was repeated four times. Cell suspensions were diluted in PBS and plated on YPD agar in duplicate. After 2 days of incubation at 30 °C, number of colony-forming-units (CFU) was quantified.

### Live-cell microscopy of neutrophil inhibition of fungal filamentation, NETosis and death

Neutrophils were seeded in 96-well plates (2 × 10^4^ cells/well) in RPMI-1640 containing 0.5 µM SYTOX Green (Invitrogen) and were allowed to settle for 15 min at 37 °C with 5% CO_2_. Neutrophils were infected by adding 50 µL (MOI:1, 2 × 10^4^ cells/well) mScarlet-expressing *C. albicans* cells (69). Neutrophils and *C. albicans* were cultured in the presence or absence of human albumin (5 mg/mL) and/or IL-1β (10 ng/mL). Wells were imaged in an Incucyte SX5 Live-Cell Analysis System (Sartorius AG). Phase contrast, green (excitation, 453–485 nm; emission, 494–533 nm) and orange (excitation: 546-568 nm; emission: 576-639 nm) images were taken every 20 minutes for 16 hours at 20× magnification. Hyphal length was quantified with the NeuroTrack analysis tool (Incucyte software 2024B; Sartorius) (70). The tool’s nuclei parameter was used to identify SYTOX Green-positive NETosis events. The analysis definition can be shared upon request. Following incubation, culture supernatants were collected for IL-1β ELISA.

### ROS production by neutrophils

Neutrophils were seeded in white 96-well plates at 5 × 10^4^ cells/well, and stimulated by addition of 50 μL *C. albicans* 1 × 10^6^ cells/mL with or without 5mg/mL albumin, or 250 ng/mL Phorbol 12-myristate 13-acetate (PMA; Sigma). Luminol-horseradish peroxidase (HRP) solution (200 µM luminol and 16 U HRP) was added (100μL) and chemiluminescence was measured every 2.5 minutes for 3 hours at 37°C in a microplate reader (Tecan M-Plex). For assays with PMA, albumin (5 mg/mL) was injected 10 min after initiation of measurements.

### ASC reporter assay

BMDMs from R26-CAG-ASC-citrine mice (39) were adjusted to a concentration of 2 × 10^5^ macrophages/mL and 300 μL of this cell suspension were seeded in an 8-well microscopy slide (ibidi), and were rested overnight. Macrophages were primed for 2 h with 50 ng/mL LPS before adding *C. albicans* SC5314 at MOI 2 in the presence (10 mg/mL) or absence of albumin. Samples were incubated (37 °C, 5% CO_2_) and fixed after 5 h using 4% formaldehyde (Carl Roth). Nuclei of macrophages were stained using DAPI (Sigma Aldrich, Merck) to determine total amount of macrophages per image. Microscopic images were acquired at a 20 × magnification using an Axio Observer Z1 (Carl Zeiss Microscopy). Number of macrophages and amount of ASC specks were determined using the counting tool in Zen (Carl Zeiss Microscopy).

### RNA isolation

VECs or hMDMs were seeded in 6-well plates and infected with *C. albicans* MOI 1 in the presence or absence of 5 mg/mL albumin. Samples for RNA isolation were collected 3 h after infection. As control, samples of *C. albicans* cells alone were taken at the same time point. RNA sample collection and isolation were done as previously described (71). Briefly, the well content was removed and replaced with 500 μL of RNeasy Lysis (RLT) buffer (Qiagen), containing 1% β-mercaptoethanol (Roth). Cells were detached using a cell scraper, immediately shock-frozen in liquid nitrogen, and stored at −80 °C until further use. Collected samples were defrosted on ice and centrifuged for 10 min (20,000 × g, 4 °C). Fungal RNA was isolated from the pellet, using a freezing-thawing method. RNA quantity and integrity was determined using a NanoDrop 1000 Spectrophotometer (Thermo Fisher Scientific) and Agilent 2100 Bioanalyzer (Agilent Technologies), respectively.

### Reverse transcription-quantitative PCR (RT-qPCR)

Isolated RNA was treated with DNase I (RNase-Free DNase Set, Qiagen) following the manufacturer’s recommendations and subsequently transcribed into cDNA using 0.5 µg Oligo(dT)12-18 Primer, 200 U Superscript™ III Reverse Transcriptase and 40 U RNaseOUT™ Recombinant RNase Inhibitor (Thermo Fischer Scientific). Obtained cDNA was used for qRT-PCR with GoTaq® qPCR Master Mix (Promega) in a CFX96 thermocycler (Bio-Rad). The expression levels were normalized against 18s rRNA. All primers used are listed in **Supplementary Table S1**.

### Fungal killing by hMDMs

Human MDMs were seeded in 96-well plates at 4 × 10^4^ cells/well and rested overnight as described above. For infection, macrophages were infected with *C. albicans* MOI 1 in the presence or absence of 10 mg/mL albumin. Samples were incubated at 37 °C and 5% CO_2_. After 3 h, the content of the wells was scraped, resuspended and transferred to a collection tube. Afterwards, sterile H_2_O was added to each well and remaining content of the wells was lysed, resuspended and transferred to corresponding collection tubes. The procedure was repeated four times. Cell suspensions were diluted in PBS and plated on YPD agar plates in duplicate. After 2 days of incubation at 30 °C, number of colony-forming-units (CFU) was quantified.

### Phagocytosis and intraphagosomal hyphal length assay

Human MDMs were adjusted to a 2 × 10^5^ cells/mL and 300 μL of this cell suspension were seeded in an 8-well microscopy slide (ibidi) and rested before the experiments as described above. Cells were infected with *C. albicans* MOI 1 and in the presence or absence of 10 mg/mL albumin. To synchronize phagocytosis, samples were centrifuged at 200 × *g* for 2 min before incubation at 37 °C and 5% CO_2_. For quantification of phagocytosis, cells were fixed after 30 min using 4% formaldehyde. For analysis of the intraphagosomal hyphae length, samples were fixed after 2.5 h. Extracellular fungal cells were stained using 50 μg/mL Concanavalin A - AlexaFluor 647 (Invitrogen, Thermo Fisher Scientific). Macrophages were washed twice using PBS and permeabilized by exposing the samples for 5 min to 0.5% Triton X (Sigma). Samples were washed with PBS again and intracellularly located *C. albicans* was visualized using 50 μg/mL calcofluor white (Fluorescent brightener 28, Sigma). Finally, samples were washed using H_2_O and stored in PBS. Microscopy was performed using an Axio Observer Z1 (Carl Zeiss Microscopy) at 20 × magnification. Image analysis was done using the counting tool and the spline curve tool in Zen software (Carl Zeiss Microscopy). All visible yeasts and hyphae per image were measured to prevent biased analysis.

### Growth curve of C. albicans in the presence of albumin and hydrogen peroxide

Growth curves of *C. albicans* SC5314 were done by seeding 1 × 10^4^ fungal cells/well in 96-well plates in RPMI-1640. Respective conditions included 5 mg/mL albumin or 1 mM H_2_O_2_ (Roth) were added. Samples were sealed with foil and incubated at 37°C in a Tecan M-Plex reader. The optical density at 600 nm was measured every 30 min for 24 h and the plate was shaken for 15 sec at 140 rpm before each measurement.

### Flow cytometry with GFP reporter strains

GFP-reporter strains were preincubated for 1 h in RPMI-1640 with and without 5 mg/mL albumin at 37 °C and 180 rpm. Subsequently, cells were washed once with RPMI-1640 and incubated for 2 h in RPMI-1640, containing 50 mM H_2_O_2_ at 37 °C and 180 rpm. Afterwards, cells were centrifuged for 5 min at 5000 × g and fixed with 4% formaldehyde for 15 min. Finally, cells were resuspended in 400 µL PBS and measurements were performed with a FACSVerse (BD Bioscience) flow cytometer. Data analysis was performed using FlowJo v.10.

### Statistical analysis

GraphPad Prism 10 was used for analyses. All data presented in the manuscript are the results of at least two independently conducted experiments. Statistical tests are indicated in the figure legends and significances are depicted in the figures: * = *P ≤* 0.05, ** = *P ≤* 0.01, *** = *P ≤* 0.001 or **** = *P ≤* 0.0001.

## Acknowledgements

We thank Julia Mantke, Maximilian Himmel, Emilia Wolf, and Candela Fernández-Fernández for technical support of *in vitro* experiments. We thank Maximilian Rothe for technical assistance and mouse breeding. We thank Wibke Krüger for her support with the Luminex measurements. We thank Ilse Jacobsen for providing access to the flow cytometry and Magpix infrastructure and Ketema Abdissa and Sigrun Kirste for advice with histology. We thank Salomé LeibundGut-Landmann for the provision of the *C. albicans* isolate 101, Julian Naglik for the provision of the *C. albicans* isolate C81, Patrick van Dijck for provision of the *C. albicans* isolate CA3153. We thank Norman Häfner, University Clinic Jena for providing SW954 cells. We thank Tim Bastian Schille (Hube lab) for sharing a fluorescent *C. albicans* strain. We thank Leo Joosten (RadboudUMC) for providing Anakinra.

## Funding

MSG, SH, and KS were supported by the German Research Foundation (Deutsche Forschungsgemeinschaft - DFG) Emmy Noether Program (project no. 434385622 / GR 5617/1-1) to MSG. KC, MSG and GGGD were supported by a Research Foundation Flanders (FWO)-funded SBO project <u>De</u>fining <u>V</u>aginal Candidiasis <u>E</u>lements of Infection and <u>R</u>emedy DeVEnIR (project number S006424N). AD was supported by an Exploration Grant of the Boehringer Ingelheim Foundation (BIS) to MSG and an ESCMID research grant 2025 to AD. This research was supported by the Free State of Thuringia and co-funded by the European Union – Project-ID 2023 FGI 0004. “A Live broadcast of the interactions between host and fungal pathogens” to MSG and AD. MP was supported by the DFG within the Collaborative Research Centre (CRC)/Transregio (TRR) 124 “Pathogenic fungi and their human host: Networks of Interaction - FungiNet” project C1 (DFG project number 210879364) to MSG. CB and SW were supported by the DFG within the CRC/TRR 241 “IECinIBD” project A03. SA was supported by funding from the European Union’s Horizon 2020 research and innovation program under grant agreement 847507 (HDM-FUN) to BH. BH was further supported by the DFG Priority Program 2225 “Exit Strategies of extracellular pathogens”. The funding providers had no role in study design; collection, analysis and interpretation of data; in the writing of the article; nor in the decision to submit the article for publication.

## Conflict of interest statement

G.G.G.D. is the chairperson of Femicare vzw and has worked as a medical consultant for various industries. None of these organizations or companies were involved in funding, the design, communication, or data analysis of this study. The other authors declare no competing interests and no conflicting financial interests.

## Data and materials availability

All data are available in the main text or the supplementary materials and will be made available upon request.

## Author contribution statement

Conceptualization: MP, TZ, MSG

Methodology: OE, DMDG, TP, EL, AD, SWe, CB, BH

Investigation: SA, AD, GV, KOC, SUJH, KSS, MJ, NJ, SWi, OE, CB, GR, VO, GGGD, MP, TZ

Visualization: SA, GV, AD, KOC, SUJH, DM, TZ, MSG

Funding acquisition: BH, GGGD, MSG

Project administration: SA, AD, MSG

Supervision: BH, MP, TZ, MSG

Writing – original draft: SA, MSG

Writing – review & editing: SA, AD, SUJH, DDG, GGGD, BH, MP, TZ, MSG

## Notes

### Competing Interest Statement

G.G.G.D. is the chairperson of Femicare vzw (www.facebook.com/profile.php?id=100063440664962) and has worked as a medical consultant for various industries. None of these organizations or companies were involved in funding, the design, communication, or data analysis of this study. The other authors declare no competing interests and no conflicting financial interests.

