## Supplement for "Local albumin excess exacerbates Candida albicans-induced inflammasome activation linked with hyperinflammation during vulvovaginal candidiasis"

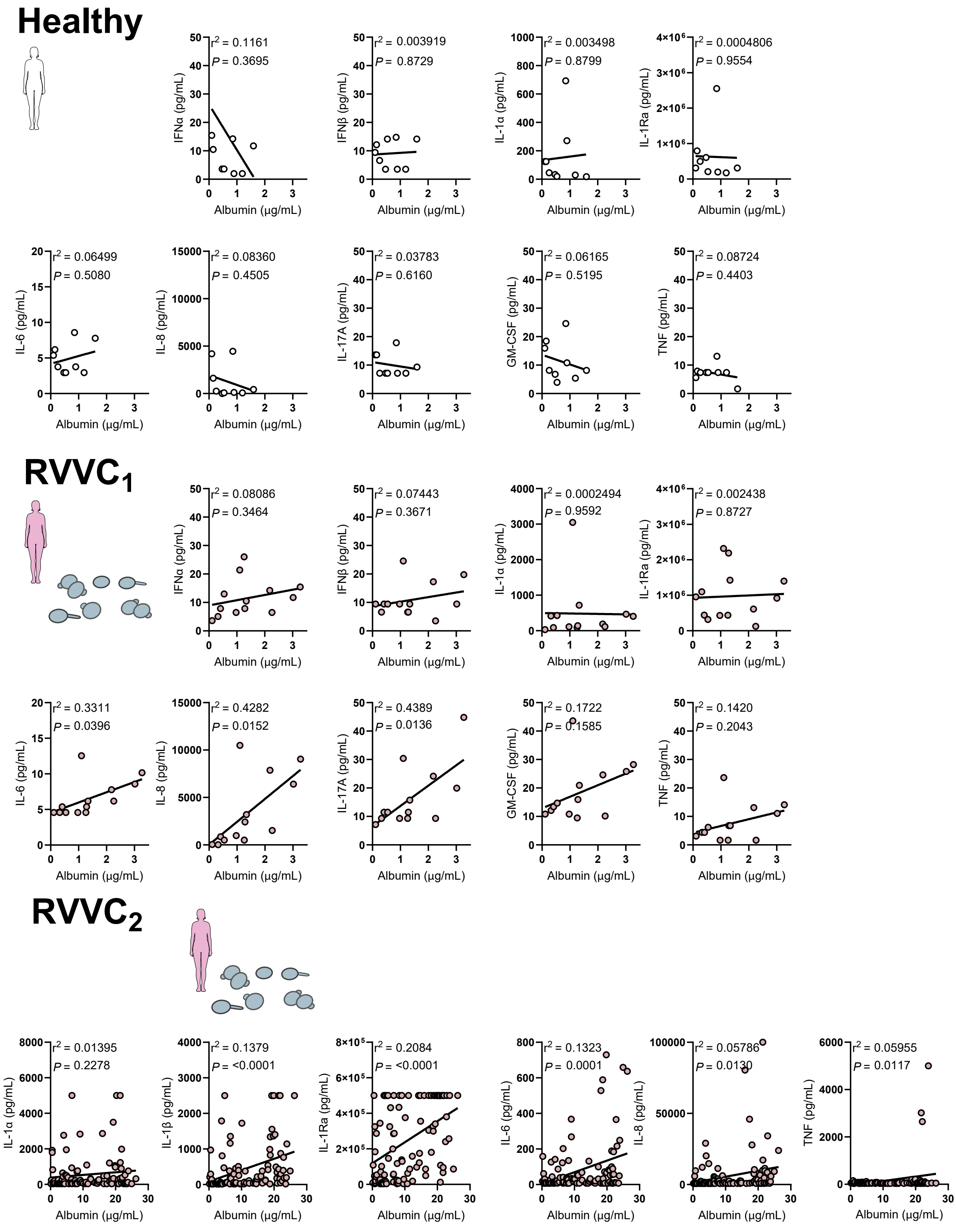


**Figure S1: Albumin concentrations correlate with VVC-associated cytokines in vaginal lavages from RVVC patients.** Cytokine and albumin concentrations were quantified in healthy women (*n* = 9) and patients from two RVVC cohorts ($RVVC_1$ *n* = 13; $RVVC_2$ *n* = 106). Correlation between cytokine levels and albumin content was assessed using Pearson correlation analysis. Pearson correlation coefficient ($r^2$) and *P*-values are indicated.

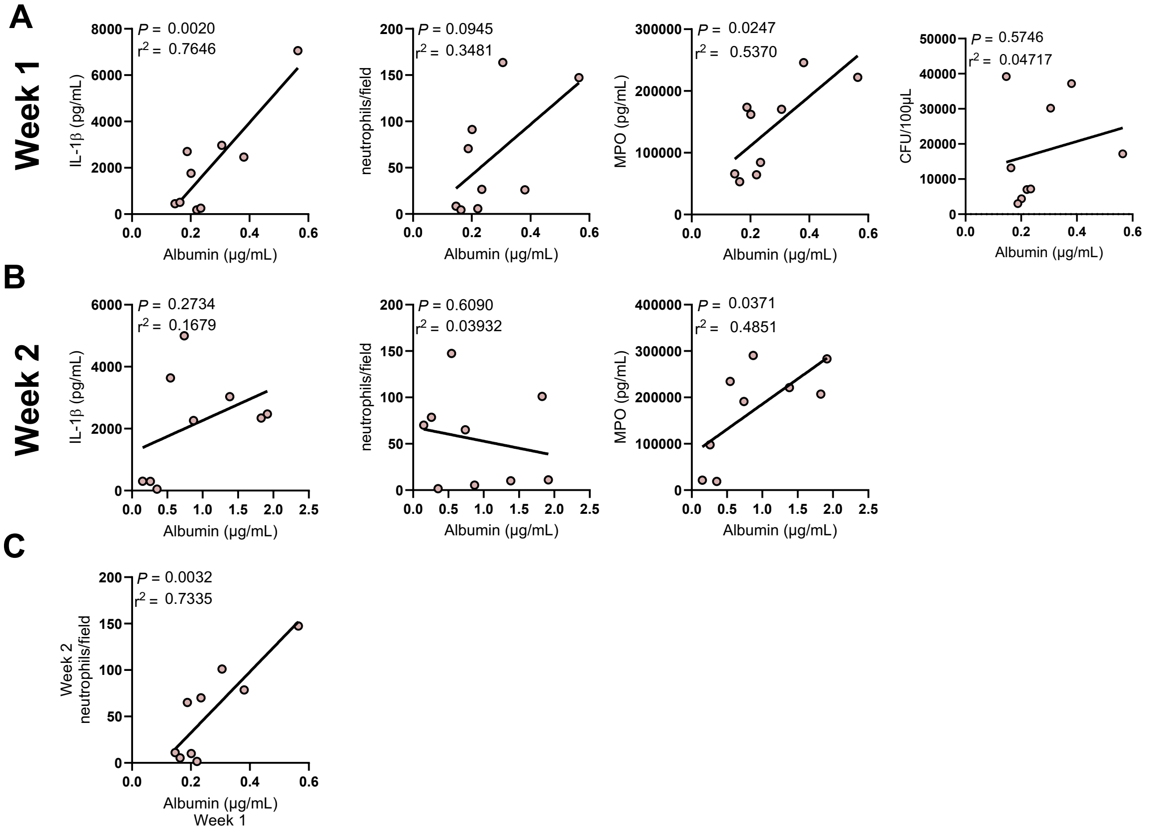


**Figure S2: Vaginal albumin concentrations correlate with VVC-associated inflammatory aspects.** (**A**,**B**) Correlations between vaginal albumin concentrations and colony forming units (CFUs; *n* = 9), IL-1β release (*n* =9), MPO release (*n* = 9) or neutrophil count (*n* = 9) in vaginal lavages (A) one week or (B) two weeks after infection of β-estradiol-treated mice with *C. albicans.* (**C**) Correlation between neutrophil count (*n* = 9) two weeks after infection of β-estradiol-treated mice with *C. albicans* with vaginal albumin concentrations measured at one week post infection. Pearson correlation coefficient ($r^2$) and *P*-values are indicated.

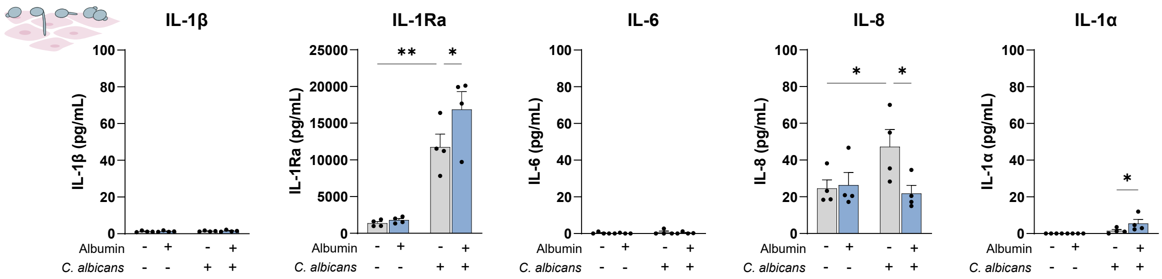


**Figure S3: Long-term epithelial infection.** Cytokine release by A-431 cells ($n$ = 5) in response to *C. albicans* infection in the presence or absence of albumin at 45 hpi . Bars represent the mean + SEM with dots as individual biological replicates. Statistical significance was determined by two-way ANOVA with Holm-Šídák post-hoc test. * = $P \leq 0.05$, ** = $P \leq 0.01$

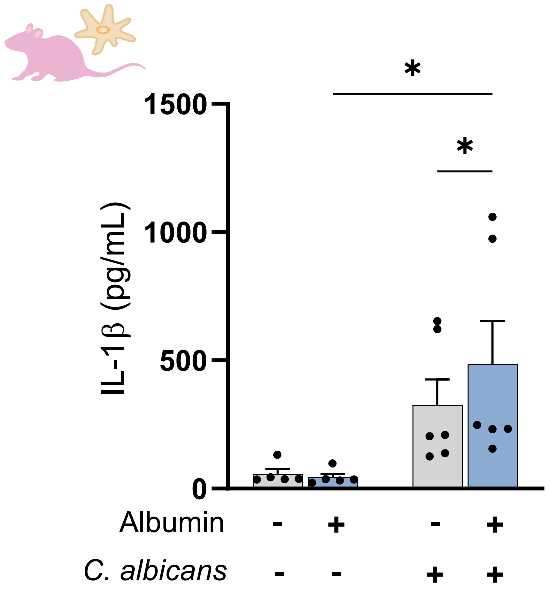


**Figure S4: Murine albumin promotes *C. albicans*-induced IL-β release in murine macrophages.** IL-1β release of murine bone-marrow derived macrophages ($n$ = 6) in response to murine albumin and *C. albicans* infection. Bars show the mean + SEM. Statistical significance was determined using a two-way ANOVA with Holm-Šídák post-hoc test. * = $P \leq 0.05$

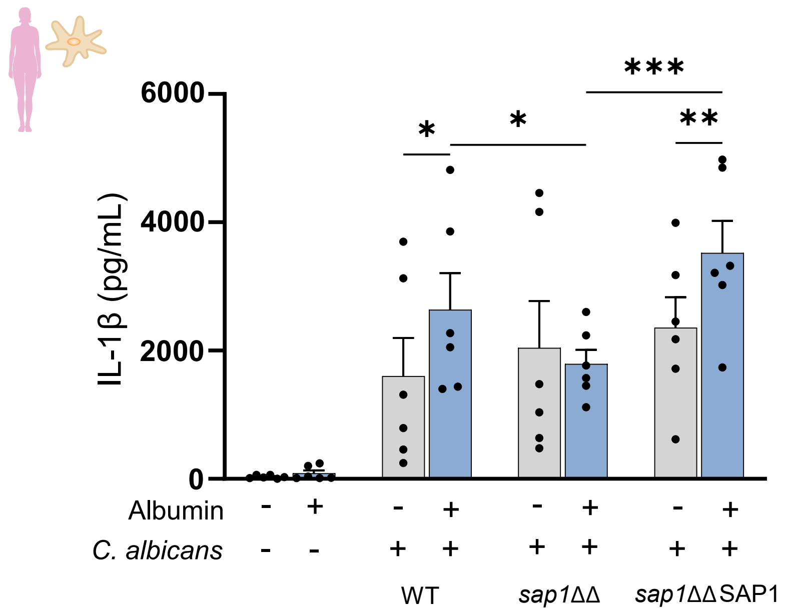


**Figure S5: Albumin-enhanced IL-1β release relies on *C. albicans* secreted aspartic proteinase 1 (Sap1).** IL-1β release by hMDMs ($n$ = 6) induced by *C. albicans* wild type, the deletion strain *sap1*ΔΔ and the revertant strain *sap1*ΔΔ *+SAP1* in the presence or absence of albumin**.** Bars represent the mean + SEM with dots as individual biological replicates. Statistical significance was determined using a two-way ANOVA including a Holm-Šídák post-hoc test. * = $P \leq 0.05$, ** = $P \leq 0.01$, *** = $P \leq 0.001$.

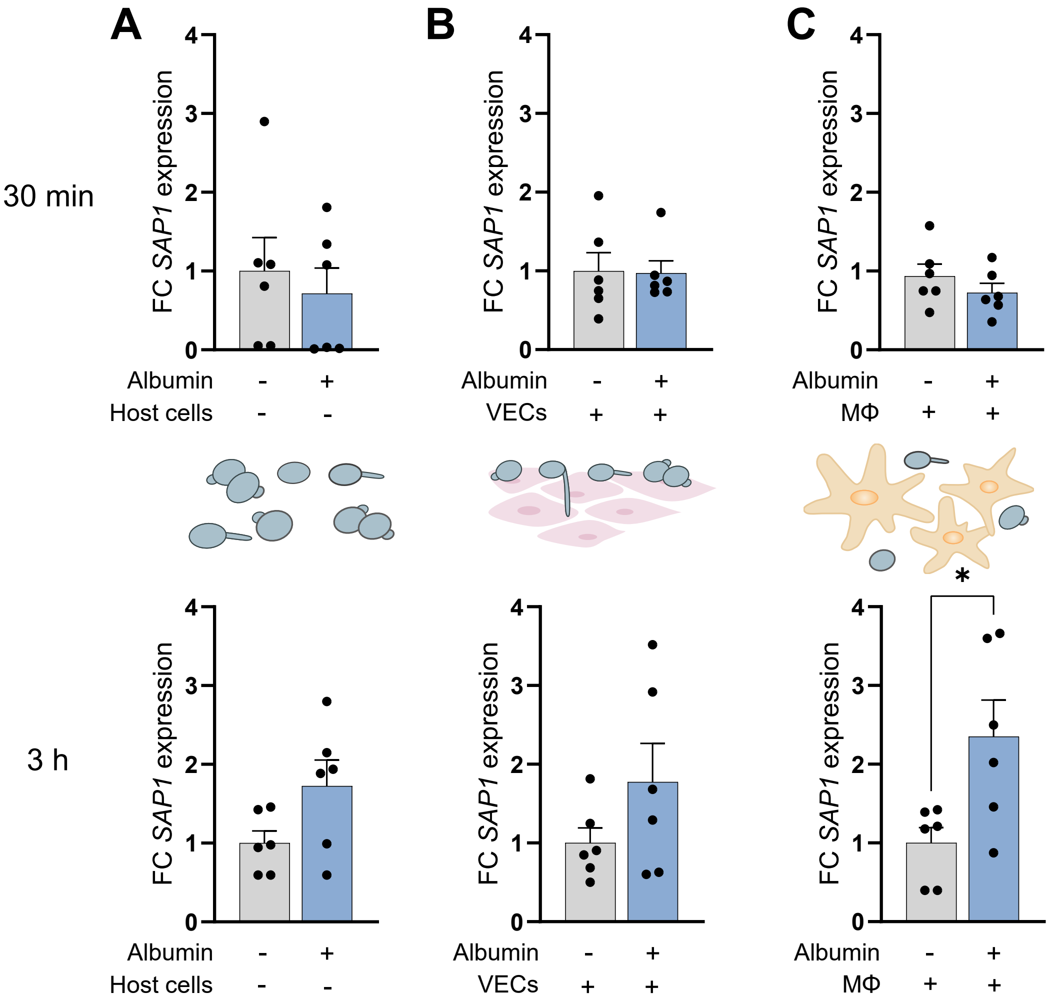


**Figure S6: mRNA expression of *C. albicans SAP1* increases in the presence of albumin.** *C. albicans* wild type was incubated on plastic (**A**), on vaginal epithelial cells (VECs; **B**) and with macrophages (MΦ; **C**). Samples for RNA isolation were collected 30 min and 3 h after infection. Gene expression was quantified by qRT-PCR. *SAP1* expression was compared to the expression of the 18S housekeeping gene. Gene expression was normalized to the mean of the control condition (without albumin) and is shown as fold change (FC). Bars show the mean + SEM and data points represent experimental replicates from individual *C. albicans* cultures (A, *n* = 6; B, *n* = 6) or individual donors (C, *n* = 6). Statistical significance was determined using a paired t-test. * = $P \leq 0.05$

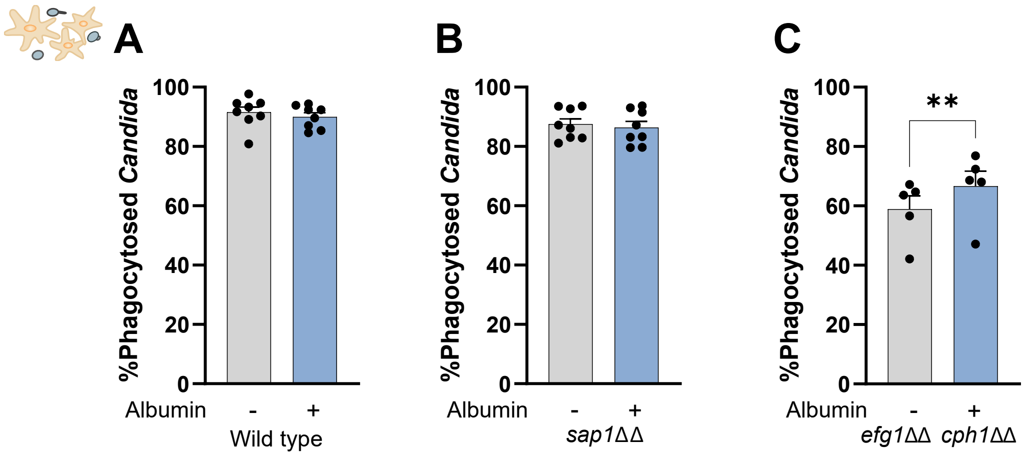


**Figure S7: Percentage phagocytosed *C. albicans* cells by hMDMs after infection in the presence or absence of albumin.** Phagocytosis of the wild type strain (**A**, *n* = 8), *sap1*ΔΔ (**B**, *n* = 8) or *efg1*ΔΔ *cph1*ΔΔ (**C**, *n* = 6) was quantified 2.5 h after infection. Bars show the mean + SEM with dots as individual biological replicates. Statistical significance was determined using a paired t-test. * = $P \leq 0.05$, ** = $P \leq 0.01$.

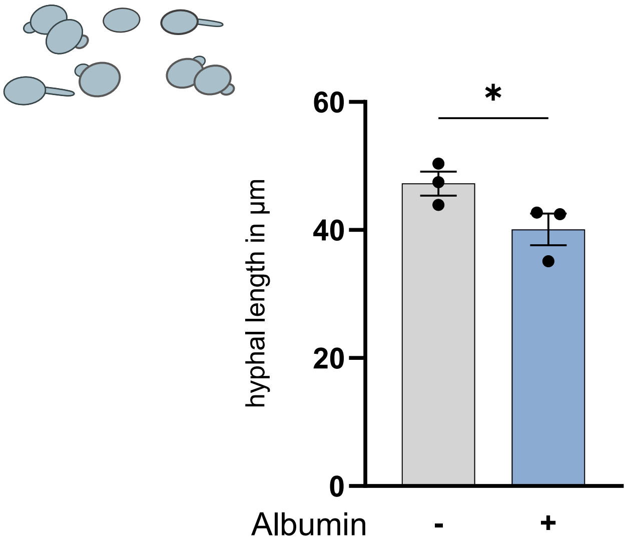


**Figure S8: Hyphal length of *C. albicans* WT in the presence or absence of albumin.** Hyphal length was quantified after 3 h growth on plastic. Bars display mean with SEM and dots are the medians of single hyphae measurements of independent experiments. Statistical significance was determined using a paired t-test. * = $P \leq 0.05$

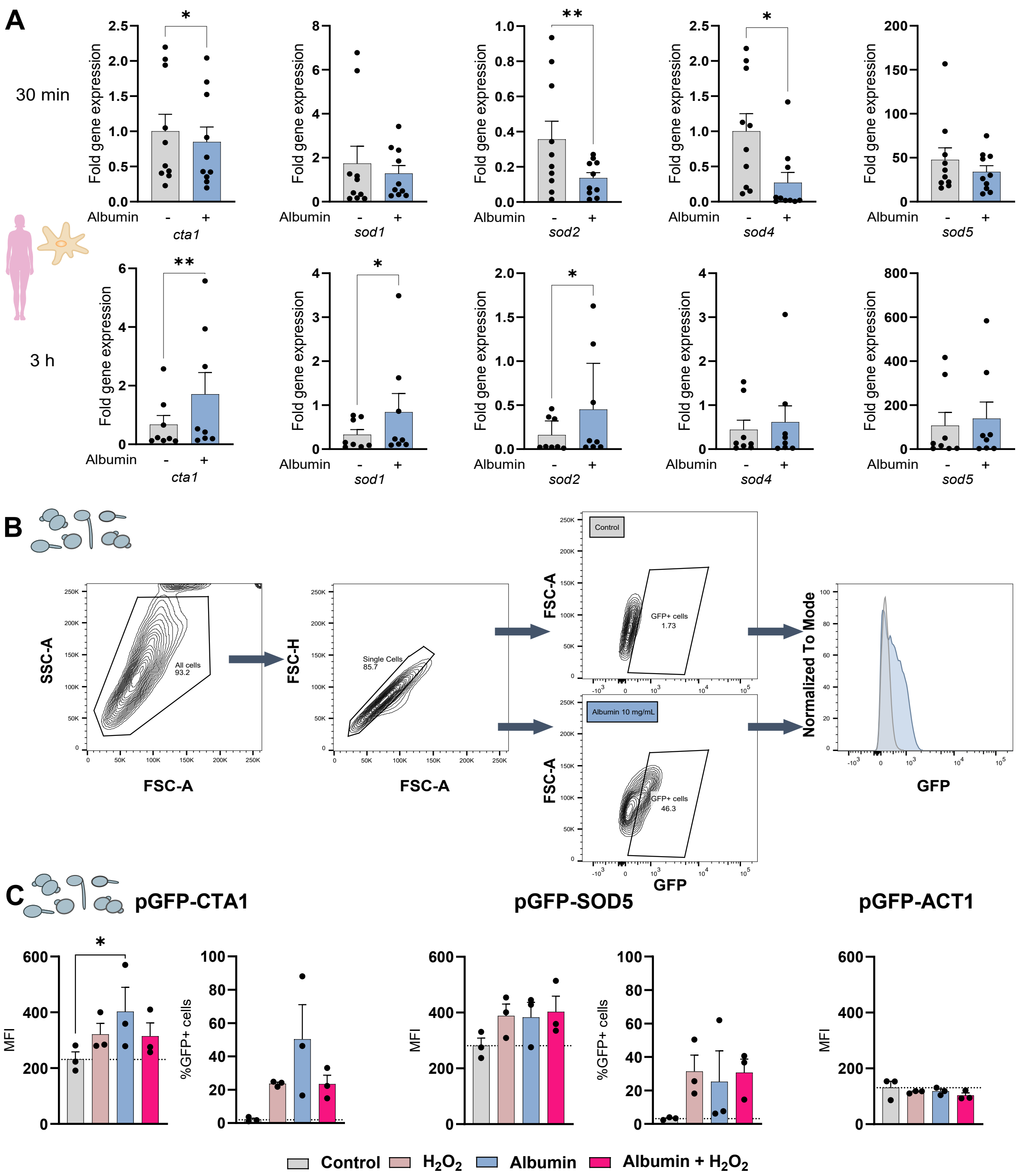


**Figure S9: *C. albicans* regulation of oxidative stress related genes in the presence of albumin. (A)** mRNA expression of oxidative stress related genes in *C. albicans* wild type during the interaction with macrophages in the presence of albumin. Samples were taken after 30 min and 3 h. Gene expression was quantified by qRT-PCR. *CTA1, SOD1, SOD2, SOD4* and *SOD5* expression was compared to the expression of the 18s housekeeping gene. Fold change (FC) gene expression normalized to the mean of the control condition (without albumin) is shown. Statistical significance was determined using a paired t-test. **(B)** Gating strategy for MFI quantification of *C. albicans* pCTA-GFP reporter strain in the presence or absence of albumin using FACS. **(C)** MFI and percentage GFP-positive cells of *C. albicans* pCTA1-GFP, *C. albicans* pSOD5-GFP and *C. albicans* pACT1-GFP. Bars show the mean + SEM with dots as individual biological replicates. Statistical significance was determined using t-test (A) or two-way ANOVA including a Holm-Šídák post-hoc test (C). * = $P \leq 0.05$, ** = $P \leq 0.01$

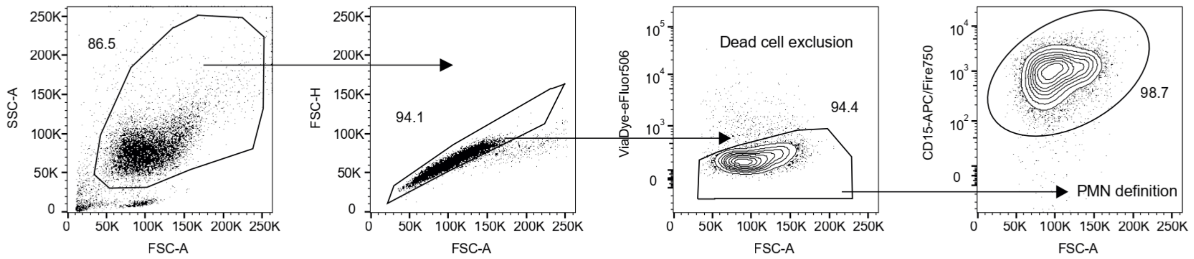


**Figure S10: Gating strategy scheme for flow cytometry of surface expression of neutrophil activation markers.** After doublet and dead cell exclusion, polymorphonuclear neutrophils (PMNs) were defined as $CD15^+$ and from this neutrophil population the mean fluorescence intensities of activation markers were extracted.

**Table S1: Primers used in this study.**

| Gene name | Primer sequence | Reference |
|---|---|---|
| *SAP1* | GATGTCATTAAAACTCCTGTTAATG<br>CCAGTTTCAATTCAGCTTGG | (1) |
| *18s* | CGATGGAAGTTTGAGGCAATA<br>CTCTCGGCCAAGGCTTATACT | (2) |
| *CTA1* | ATTTCATCCACACCCAAAGAGA<br>CAAGTAATCCCAAAACATGTTAGCA | (3) |
| *SOD1* | TGTTGTCAGAGGTGATTCAAAAGTC<br>GTTGGAGCGGATTCGGATT | (3) |
| *SOD2* | TTGGCTCCTGTCTCTCAAGG<br>CCAATTTGCCATTGGTGATT | (2) |
| *SOD4* | TTTGAGCCAGCAAACAATGG<br>CACCTGAAGGCAATCCAGTTAAA | (3) |
| *SOD5* | AAGGATTGCCCTCTGATATTGG<br>GATGCTGGCACTGGTTTTTCA | (3) |
